# Baseline cellular state dictates the molecular impact of *KRAS* mutant variants in pancreatic cancer cells

**DOI:** 10.64898/2026.03.10.710185

**Authors:** Yanixa Quiñones-Avilés, Barbora Salovska, Cassandra S. Markham, Yi Di, Benjamin E. Turk, Yansheng Liu, Mandar Deepak Muzumdar

## Abstract

*KRAS* is mutated in over 90% of pancreatic ductal adenocarcinomas (PDAC), where hotspot alterations in codons 12, 13, and 61 drive tumor initiation and progression. Although distinct biochemical properties have been described for individual KRAS mutants, whether they generate unique allele-specific signaling programs in PDAC cells remains unresolved. Here, we systematically interrogated the molecular consequences of seven common KRAS mutant variants in reconstituted isogenic, KRAS-deficient PDAC cell lines by integrated transcriptomic, proteomic, and phosphoproteomic profiling. We found that baseline cellular state, rather than allele identity, was the predominant driver of molecular variation. Comparisons with established KRAS reference signatures revealed significant but moderate overlap at the mRNA level and less so at the proteome level. Pathway analyses highlighted interferon response and mitochondrial translation as recurrently altered across alleles, while phosphoproteomic data confirmed robust ERK1/2 activity and suppression of DYRK kinase substrates by mutant KRAS expression. Importantly, no robust allele-specific molecular programs were identified. Together, our study establishes a comprehensive multi-omics resource for KRAS signaling in PDAC and demonstrates that cellular context exerts a stronger influence than allele identity in shaping molecular profiles, with implications for interpreting putative allele-specific signaling dependencies and therapeutic vulnerabilities.

## INTRODUCTION

Pancreatic ductal adenocarcinoma (PDAC) is the 3^rd^ leading cause of cancer-related death in the United States and is predicted to become the 2^nd^ by 2030 (*1*). *KRAS* mutations are considered the initiating event in PDAC development, with activating *KRAS* mutations being found in over 90% of human PDAC tumors (*2*). Single-point mutations in amino acids G12, G13, and Q61 disrupt KRAS inactivation by GTPase-activating proteins (GAPs), rendering KRAS near constitutively active and facilitating tumorigenesis (*3*). While these hotspot residues can harbor multiple substitutions, the most common alleles observed in PDAC are *KRAS^G12D^*, *KRAS^G12V^*, and *KRAS^G12R^*, together accounting for nearly 90% of cases (*2*).

Mutant KRAS proteins reportedly possess unique biochemical properties, such as distinct intrinsic and GAP–stimulated GTP hydrolysis rates and binding affinities to the RAS-binding domain of effector RAF kinases (*4*), supporting the potential for divergent allele-specific signaling networks that underscore unique vulnerabilities and response to treatment. Indeed, previous studies comparing the phosphoproteomic landscape of KRAS^G12D^ and KRAS^G13D^ mutants in isogenic colorectal cancer cell lines have reported mutant-specific proteomic and phosphoproteomic signatures (*5, 6*). In addition to these biochemical and signaling differences, biological distinctions among KRAS mutants have been reported. For example, unlike KRAS^G12D^ and KRAS^G12V^, KRAS^G12R^ cannot stimulate macropinocytosis due to its impaired ability to engage the catalytic p110α subunit of phosphoinositide 3-kinase (PI3K), suggesting allele-specific biology (*7*). Consistent with these observations, a comparison of common and uncommon KRAS mutants in isogenic MCF10A mammary epithelial cells showed that distinct alleles had differential effects on EGF-independent proliferation and anchorage-independent growth (*8*). While these studies highlight important KRAS mutant-specific biochemical and biological differences, they have largely focused on discrete comparisons of a small subset of alleles in non-pancreatic or non-malignant systems, underscoring the need to comprehensively interrogate the molecular programs engaged by a broader panel of *KRAS* mutations in relevant PDAC models.

To explore the molecular impact of different *KRAS* alleles, we generated a panel of four KRAS-deficient human PDAC cell lines using clustered regularly interspaced short palindromic repeats (CRISPR)/Cas9-mediated gene editing and established isogenic cells reconstituted to express wild-type or seven common mutant KRAS variants. We combined RNA-sequencing (RNA-Seq) and data-independent acquisition mass spectrometry (DIA-MS) to profile transcriptomic, proteomic, and phosphoproteomic changes driven by KRAS variants under steady-state conditions. Interestingly, we found that baseline cellular state, rather than the specific mutation, largely dictated the global impact of KRAS variants. Convergent transcriptomic and proteomic pan-mutant KRAS signatures reveal that mutant KRAS regulates interferon signaling response, mitochondrial translation, ERK1/2, and DYRK signaling pathways, partially overlapping with recently reported KRAS mutant signatures derived from other cell lines or cancer types (*9–11*). Taken together, our study establishes a comprehensive multi-omics resource for defining and comparing KRAS mutant signatures in PDAC. Further, our findings suggest that baseline cellular states and signaling context exert a stronger influence than allele identity, with important implications for interpreting putative allele-specific signaling and dependencies in PDAC and potentially other *KRAS* mutant cancers.

## RESULTS

### Derivation of isogenic cell models to study mutant KRAS variants in PDAC

We previously demonstrated that a subset of PDAC cells can rewire signaling to survive KRAS ablation (*12*). Importantly, these KRAS-independent cells demonstrated phenotypic reversibility in morphology, growth kinetics, and signaling when mutant KRAS was re-expressed, providing a tractable and genetically controlled platform to dissect the molecular consequences of specific *KRAS* mutations systematically in PDAC cells (*12*). We employed a similar CRISPR/Cas9-based gene-editing strategy to ablate *KRAS* expression in two human PDAC cell lines (8988T and KP4). Single cell subclones were expanded and screened for successful *KRAS* ablation by immunoblotting (**Fig. S1A-B**). Next, we selected two KRAS-deficient clones per parental cell line (8988T K275 and K328 and KP4 K22 and K63) (**Fig. S1C**). Whole exome-sequencing did not identify recurrent single nucleotide variants (SNVs) nor copy number variants (CNVs) in cancer drivers that would be predicted to endow KRAS independence (**Table S1-S3**), suggesting that these cell lines are suitable for isogenic comparisons of mutant KRAS variants. Nonetheless, the clones differed in baseline measures of canonical downstream signaling pathways: mitogen-activate protein kinase (MAPK; pERK) and PI3K (pAKT) (**Fig. S1C-D**). We reconstituted these clones with wild-type (WT) or seven common mutant *KRAS* variants co-expressed with a green fluorescent protein (GFP) reporter by lentiviral transduction, selecting for similar KRAS expression by fluorescent-activated cell sorting (FACS) (**Fig. 1A**). Importantly, we confirmed comparable KRAS expression levels by immunoblotting (**Fig. 1B**). This controlled expression enabled direct comparisons across KRAS mutants within each cellular background, ensuring that observed differences reflect mutant-specific biology rather than variability in protein expression.

**Figure 1.**
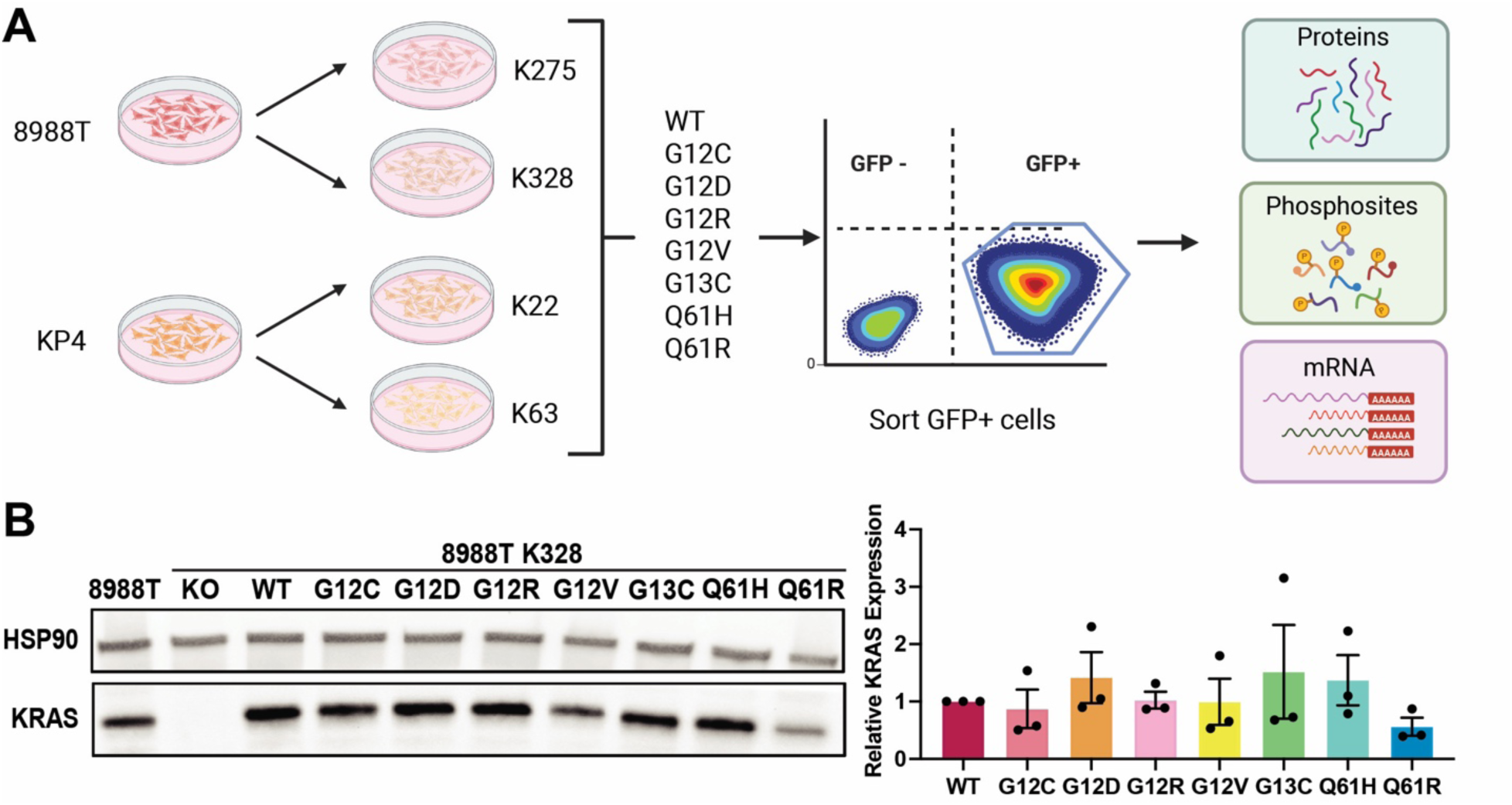
**Generation and molecular profiling of isogenic *KRAS* mutant PDAC cells.** (A) Experimental workflow of multi-omics profiling (RNA-seq, DIA-MS proteome, and phosphoproteome) of two *KRAS* knockout clones derived from each parental PDAC cell line (8988T and KP4) reconstituted with wild-type (WT) or seven different KRAS mutants to assess KRAS-dependent molecular changes. KRAS was co-expressed with GFP, which was used for sorting cells of similar fluorescence. Generated using Biorender.com. (B) Representative immunoblot and quantification of relative KRAS protein expression (mean ± SEM of n = 3 biological replicates) of each mutant (vs. WT, normalized to loading control HSP90) showing comparable KRAS expression across reconstituted 8988T K328 cells. KRAS expression was not significantly different to WT (repeated measures one-way ANOVA with Dunnett’s post-hoc test).

### Multi-omic profiling of KRAS mutations in isogenic human PDAC cell lines

To determine the molecular effects of mutant KRAS expression in PDAC cells, we quantified the steady-state transcriptome, proteome, and phosphoproteome across all four isogenic cell line series via RNA-Seq and DIA-MS-based proteomics analyses. Multi-omics profiling confirmed deep coverage of the transcriptome, proteome, and phosphoproteome: RNA-Seq identified, on average, transcripts of over 14,000 protein-coding genes, while DIA-MS identified more than 7,500 unique protein groups across all samples (**Fig. S2A-B**). Similarly, between 35,000-40,000 unique class-I phosphosites (*13*) were detected across all samples, with the exception of K22 KRAS^G13C^, which yielded ∼17,000 sites (**Fig. S2C)**. The lower phosphosite coverage in this sample likely resulted from a small sample input, as a repeat biologic replicate identified ∼37,000 sites but showed a batch effect during data integration (**Fig. S2C-D**). As a result, this sample was excluded from downstream phosphoproteomic analyses.

We performed several assessments to validate the quality of the multi-omic data for mutant KRAS comparisons. First, KRAS expression was comparable across all samples in the proteomics data (**Fig. S3A**), consistent with immunoblotting results (**Fig. 1B**). Second, the average absolute correlation between the replicates in the proteome and phosphoproteome dataset was ρ=0.99 and ρ=0.93, respectively, demonstrating excellent technical reproducibility (**Fig. S3B**). Third, we assessed concordance between omic layers. Pairwise Spearman correlation analysis revealed a moderate positive relationship (ρ=0.45-0.65 across samples) between mRNA and protein expression changes across KRAS variants (**Fig. S3C**), indicating that a considerable proportion of proteomic variation mirrors transcriptomic changes in steady-state conditions. Similar mRNA-protein level correlations have been observed widely in other studies independent of the cell lines analyzed (*14*). Protein-phosphosite correlations also revealed a moderate positive relationship between both layers (ρ=0.45-0.61) (**Fig. S3C**), consistent with previous reports (*15*). Therefore, these data provide a rigorous, high-quality, and comprehensive molecular resource to study KRAS mutant- and allele-specific pathway engagement in PDAC cells.

#### Defining molecular signatures of KRAS mutation

To evaluate how mutant KRAS (KRAS^MUT^) variants differentially affect PDAC cells, we first performed unsupervised hierarchical clustering for each independent omics layer (**Fig. 2A**). Although mutant KRAS-expressing cells largely clustered away from wild-type KRAS (KRAS^WT^) within each cell line, there were no consistent global clustering patterns for specific mutant alleles across the mRNA, proteome, and phosphoproteome layers. Furthermore, cell line identity, rather than mutant KRAS expression, was the primary driver of molecular variability across samples and omics layers, even separating clones derived from the same parental background (**Fig. 2A**). Interestingly, differential expression analyses revealed a spectrum of responsiveness to KRAS^MUT^ reconstitution across cell lines. 8988T K275 displayed the most pronounced transcriptomic, proteomic, and phosphoproteomic changes, while 8988T K328 showed minimal changes and KP4-derived clones exhibited an intermediate response (**Fig. 2B**). We observed a similar pattern of cell-specific responsiveness in PI3K and especially MAPK signaling when analyzed by immunoblotting (**Fig. 2C**), arguing that these canonical pathways may, in part, govern the differential impacts of KRAS^MUT^ expression on individual cell lines. Together, these findings suggest that the baseline cellular state of the clones strongly influences the impact of *KRAS^MUT^* allele expression on the cellular transcriptome, proteome, and phosphoproteome.

**Figure 2.**
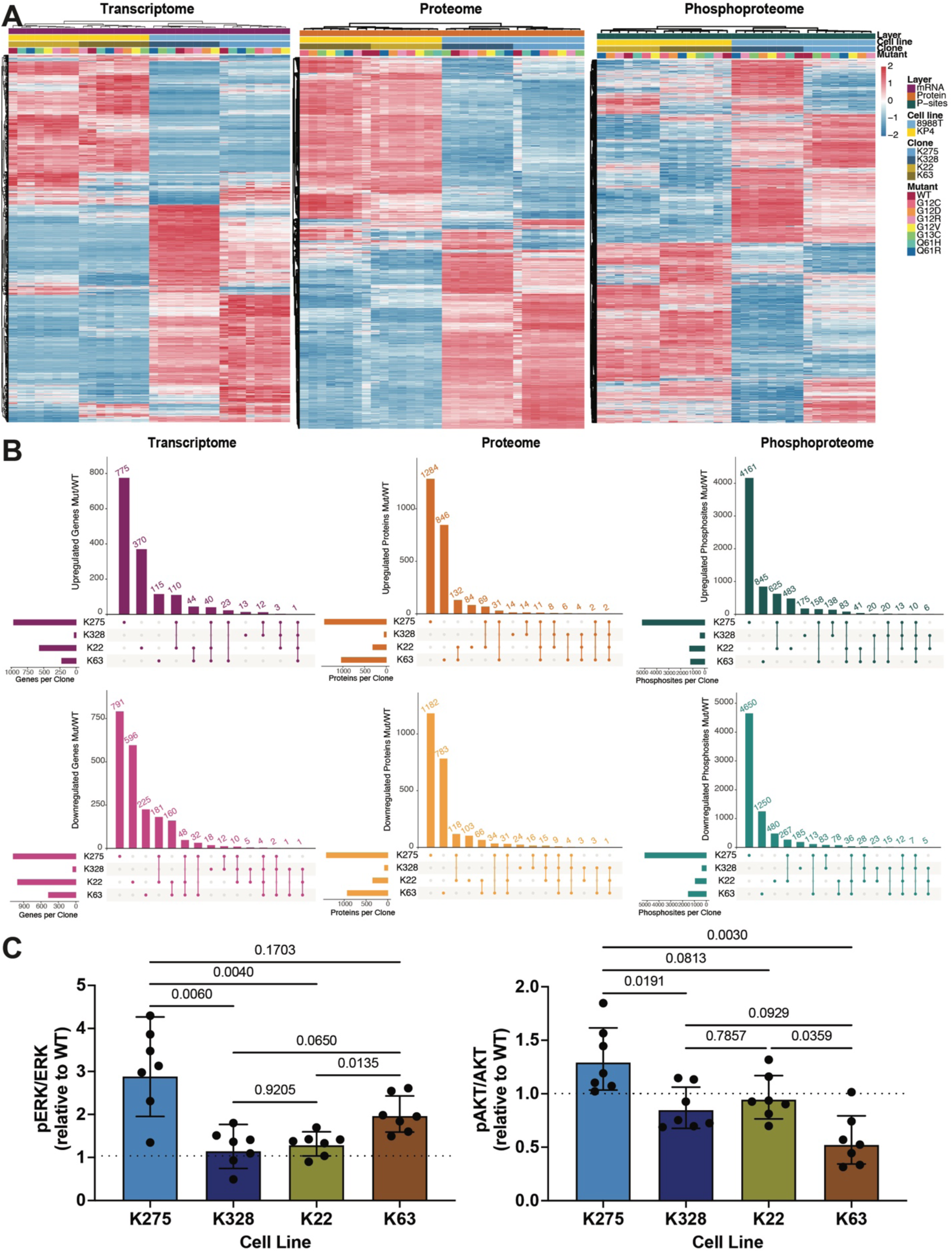
**Baseline cellular state dictates the impact of mutant KRAS expression.** (A) Unsupervised hierarchical clustering of transcriptomic, proteomic, and phosphoproteomic datasets show that clone origin, not *KRAS* allele, is the major driver of sample segregation, even between clones derived from the same parental line. Color scale denotes row-normalized absolute abundance. (B) UpSet plots showing the overlap of significantly upregulated (top) and downregulated (bottom) genes (transcriptome), proteins (proteome), and phosphosites (phosphoproteome) across all four reconstituted cell lines. A fold change ≥ 1.3 and adjusted *p*-value ≤ 0.05 threshold were used to determine differential expression of all KRAS^MUT^ relative to KRAS^WT^ for at least two cell lines. Each bar represents the number of shared or unique differentially expressed features between clones. Shared upregulated and downregulated features highlight the limited global convergence of KRAS-dependent molecular responses across cell lines. (C) Quantification of pERK1/2 (T202/Y204) and pAKT (S473) levels relative to total ERK and AKT levels in each cell line. The ratios (geometric means ± geometric SD) of the expression KRAS^MUT^ (n = 7 mutants) relative to KRAS^WT^ (average of n = 3 biologic replicates) for each cell lines are shown. *p*-values are derived from Brown-Forsythe lognormal ANOVA with Games-Howell’s post-hoc test.

We next derived a steady-state KRAS^MUT^ signature by selecting features consistently differentially expressed comparing all KRAS^MUT^ variants relative to KRAS^WT^ in at least two of the four cell lines, thereby enriching for changes that are not clone-specific but reflect shared KRAS^MUT^-dependent programs. Using a false discovery rate (FDR) threshold of 0.05 and an absolute fold-change cutoff of 1.3, the resultant KRAS^MUT^ transcriptomic signature was comprised of 233 upregulated and 456 downregulated genes (**Table S4**). Similarly, differential expression analysis of proteomic data defined a KRAS^MUT^ signature of 277 upregulated and 298 downregulated proteins, mirroring the extent of reprogramming observed at the transcriptomic level (**Table S4**). At the phosphoproteome level, we identified 1,050 upregulated and 621 downregulated phosphosites, assigned to 531 and 298 distinct phosphorylated proteins, respectively (**Table S4**).

We next performed enrichment analysis to identify hallmark pathways, processes, and kinases systematically altered by KRAS^MUT^ expression across the transcriptome, proteome, and phosphoproteome. Across functional pathways, we observed substantial overlap at the mRNA and protein levels, with notable differences in directionality and strength of enrichment (**Fig. 3A** and **S4**). The strongest convergent association we observed for both the transcriptome and proteome across functional databases was downregulation of interferon signaling response (“interferon alpha” and “gamma response” in the Molecular Signatures Database (mSigDB) Hallmark gene sets, “interferon signaling” and “interferon alpha/beta signaling” in Reactome Pathways, and “negative regulation of viral genome replication” in Gene Ontology (GO): Biological Processes) with KRAS^MUT^ (**Fig. 3A** and **S4)**. Strikingly, mitochondrial translation pathways (GO: Biological Processes and Reactome Pathways) were strongly and uniquely enriched in the upregulated proteome (**Figure S4**). To gain insights into additional signaling events altered in KRAS^MUT^ PDAC cell lines, we performed kinase motif enrichment analysis (KMEA) (*16*) using the KRAS^MUT^ phosphoproteomic signature. We observed strong enrichment for upregulated sites conforming to the ERK1/2 motif as well as motifs for other related MAPKs (JNK and p38) and cyclin-dependent kinases (CDKs) (**Fig. 3B**). Conversely, we observed a distinct dual-specificity tyrosine phosphorylation-regulated kinases (DYRKs) signature among the downregulated sites (**Fig. 3B**). Together, our multi-omics KRAS^MUT^ signatures provide a resource for defining the convergent effects of KRAS^MUT^ expression in PDAC cells.

**Figure 3.**
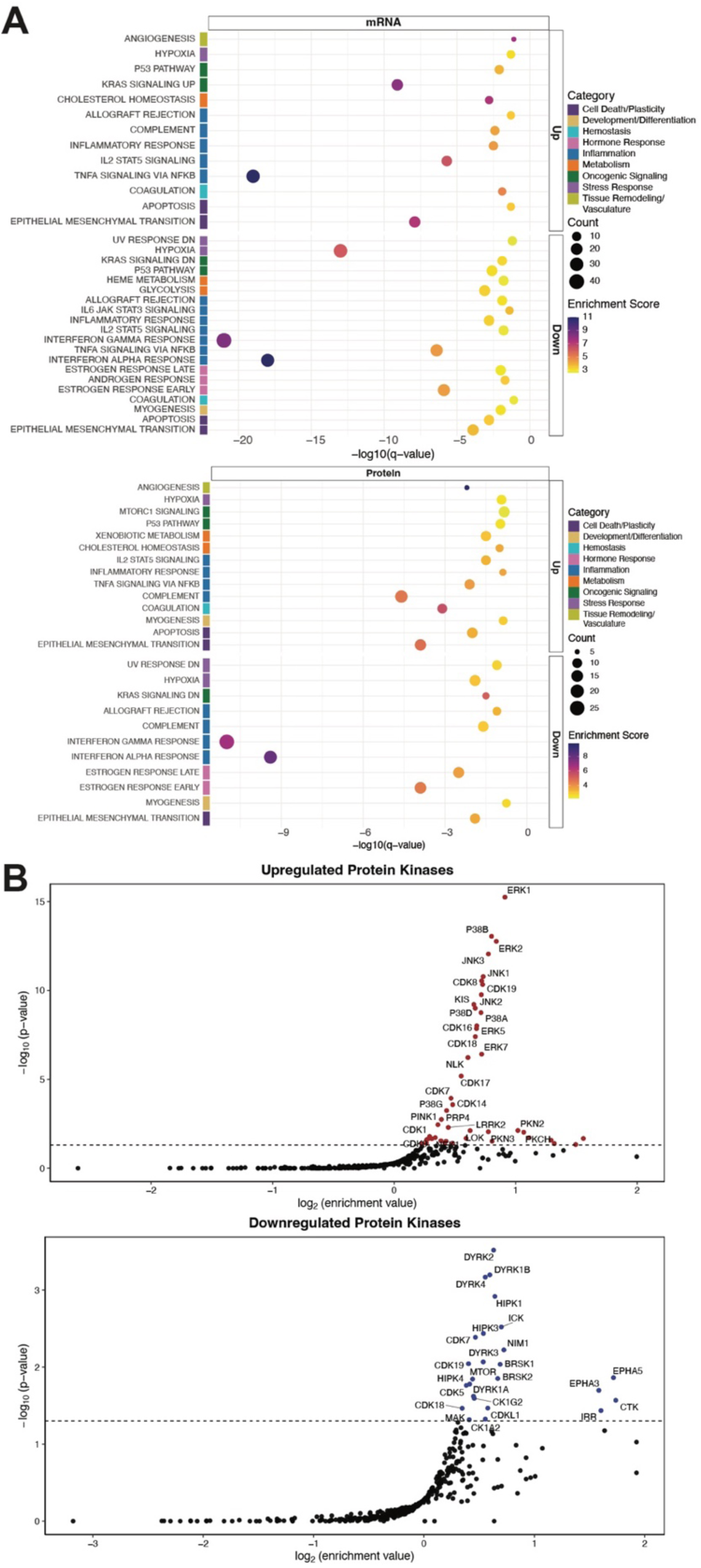
**Functional pathways and kinase activity associated with mutant KRAS expression.** (A) Pathway-level enrichment using the mSigDB Hallmarks dataset for the transcriptomic (mRNA) and proteomic (protein) differences (upregulated (Up) or downregulated (Down)) for the KRAS^MUT^ multi-omic signatures defined by selective differentially expressed hits (FC ≥ 1.3 and FDR ≤ 0.05) in at least 2 out of 4 clones (≥2 clones) compared to KRAS^WT^. Color scale denotes enrichment score. Circle size designates gene count overlap. −log_10_(*q*-value) of hypergeometric test is plotted. (B) Kinase motif enrichment analysis (KMEA) using phosphosites in the KRAS^MUT^ multi-omic signature reveals increased phosphorylation of ERK1/2 kinase substrates (upregulated protein kinases) and decreased phosphorylation DYRK kinase substrates (downregulated protein kinases). Log_2_ enrichment vs. −log_10_(*p*-value) is plotted.

### Towards a unified multi-omic mutant KRAS signature across datasets

The suppression of interferon response signaling and inferred enrichment of increased ERK1/2 activity and reduced DYRK activity mirrored findings from recent studies on the ERK-dependent transcriptome and phosphoproteome (*9, 10*). Therefore, we sought to rigorously compare our KRAS^MUT^ transcriptomic signature against publicly available KRAS-dependent signatures to assess concordance and biological relevance. As a benchmark, we used the widely adopted Hallmark KRAS Signaling gene sets from the MSigDB, a reference standard for defining KRAS pathway engagement across cancers (*17*). To quantify enrichment, we performed over-representation analysis of the selected gene sets, stratifying by direction of regulation (UP or DOWN). When comparing our upregulated KRAS^MUT^ transcriptomic signature with the Hallmark KRAS Signaling UP signature, we observed significant overlap (enrichment=8.48, p=1.09×10^−12^; **Fig. 4A**). Likewise, our downregulated KRAS^MUT^ transcriptomic signature also showed significant enrichment with the Hallmark KRAS Signaling DOWN gene set (enrichment=3.37, p=4.39×10^−4^; **Fig. 4A**). Despite this enrichment, only 19 and 11 specific genes overlapped between ours and the Hallmark UP and DOWN gene sets, respectively (**Table S5**). We reasoned that this may be due to the broad, pan-cancer derivation of the Hallmark gene sets using immortalized lung, breast, kidney, and prostate epithelial cells (*17*), whereas our signature captures PDAC-specific transcriptional programs induced by KRAS^MUT^. Therefore, we compared our results with the KRAS-dependent transcriptome reported by Klomp et al. (*9*) and derived from eight human *KRAS* mutant PDAC cell lines following a 24-hr *KRAS* siRNA treatment. Despite significant enrichment between our upregulated (enrichment=5.64, p=6.41×10^−23^; **Fig. 4A**) and downregulated (enrichment=3.37, p=2.07×10^−19^; **Fig. 4A**) KRAS^MUT^ transcriptomes, the number of overlapping genes (48 and 79; **Table S5**) remained modest as compared to the total number of significantly altered mRNAs we detected. The partial overlap may be explained by differences in experimental design, as our dataset represents steady-state molecular profiles, whereas Klomp et al. measured acute transcriptional responses to KRAS suppression. Importantly, only a small number of genes (n=12) overlapped across all datasets (*CELSR2*, *DUSP6*, *ETV4*, *ETV5*, *GPRC5C*, *HBEGF*, *LIF*, *MALL*, *MX1*, *SPRY2*, *SYNPO*, *TMEM158*; **Table S5**), including many genes previously described as being transcriptionally regulated by MAPK signaling (*18*). Given this sparse overlap, cellular context largely overshadows the effect of *KRAS* mutation across cell lines.

**Figure 4.**
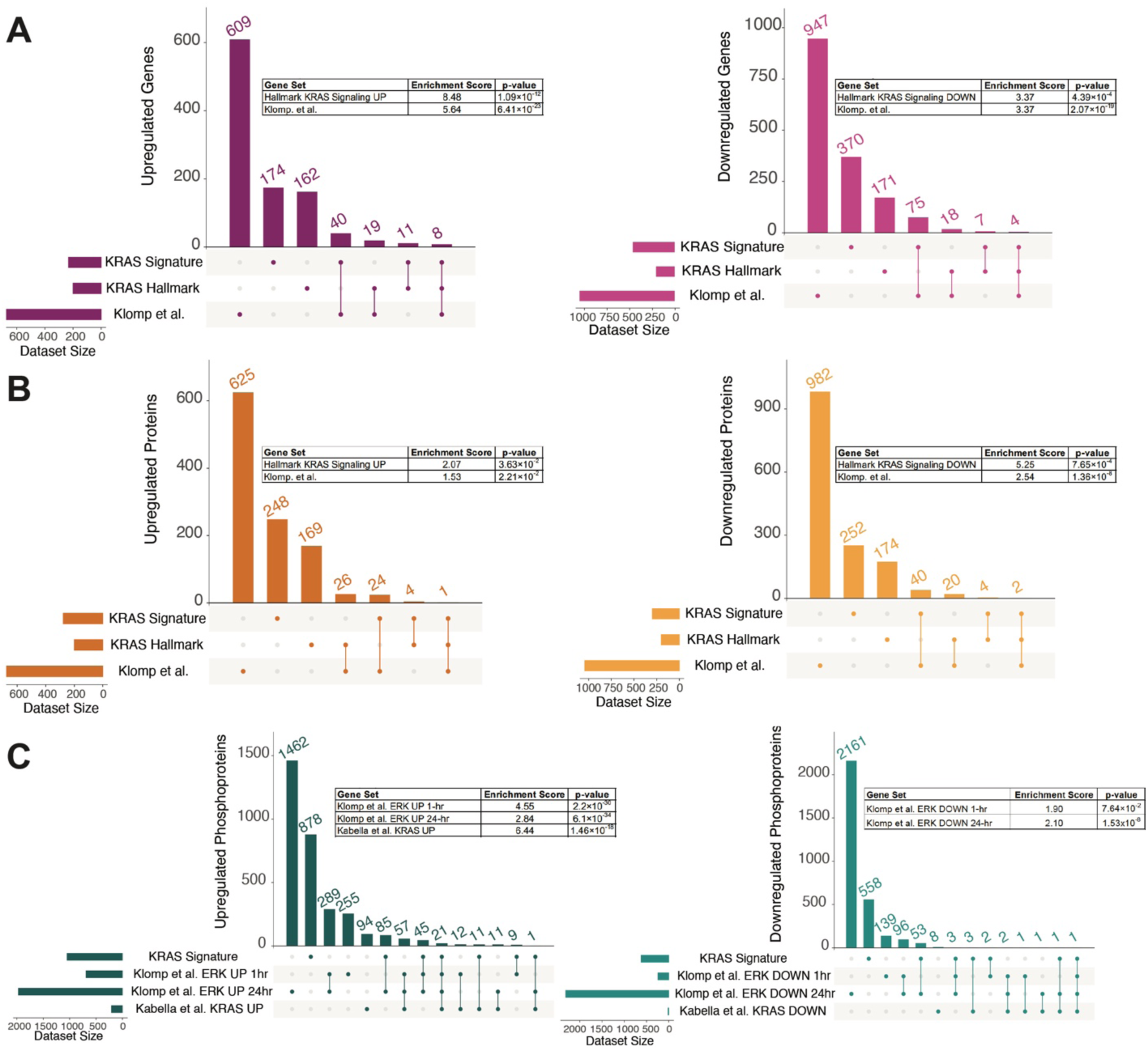
**The KRAS^MUT^ multi-omic signatures significantly overlap with prior KRAS-dependent datasets.** (A) UpSet plot illustrating overlap between the transcriptomic KRAS^MUT^ signature and previously published transcriptomic KRAS-dependent signatures (Hallmark KRAS Signaling (*17*) and Klomp *et al.*(*9*)) identifies a small core of shared KRAS-regulated genes. Enrichment scores and *p*-values of hypergeometric test are shown for corresponding upregulated or downregulated gene sets. (B) UpSet plot illustrating overlap between the proteomic KRAS^MUT^ signature proteomic and previously published KRAS-dependent transcriptomic signatures shows moderate overlap with KRAS-regulated datasets (Hallmark KRAS Signaling (*17*) and Klomp *et al.*(*9*)) Enrichment scores and *p*-values of hypergeometric test are shown for corresponding upregulated or downregulated gene sets. (C) UpSet plot illustrating overlap between the phosphoproteomic KRAS^MUT^ signature strong enrichment despite limited share sites with 1-hr and 24-hr ERK-regulated phosphoproteome and KRAS phosphoproteome core signature (Klomp *et al.*(*10*) and Kabella *et al.*(*11*)). Enrichment scores and *p*-values of hypergeometric test are shown for corresponding upregulated or downregulated gene sets.

We then compared our KRAS^MUT^ proteomic signature against the Hallmark KRAS Signaling and Klomp et al. datasets (**Fig. 4B**). The upregulated KRAS^MUT^ proteome also showed significant enrichment with Hallmark KRAS Signaling UP (enrichment=2.07, p=3.63×10^−2^), while the downregulated proteome exhibited stronger enrichment with the corresponding DOWN gene set (enrichment= 5.25, p=7.65×10^−4^; **Fig. 4B**). Comparison with the Klomp et al. signatures yielded more restricted overlap, with the upregulated proteome showing a weaker enrichment (enrichment=1.53, p=2.21×10^−2^) and the downregulated proteome displaying more significant enrichment (enrichment=2.54, p=1.36×10^−8^; **Fig. 4B**). The lower overlap at the proteomic level is not unexpected, given that both reference datasets were generated from transcriptomic data and that there is moderate correlation between mRNA and protein abundance (**Fig. S2C**) (*19*). Furthermore, only two genes (*SYNPO* and *MX1*) were consistently downregulated at both the transcriptomic and proteomic layers across all datasets (**Table S5**).

Lastly, we compared our KRAS^MUT^ phosphoproteomic signature against two recently reported reference phosphoproteomics datasets (**Fig. 4C)**: (1) the ERK-regulated phosphoproteome identified by Klomp et al. (*10*) following acute (1-hr) and prolonged (24-hr) ERK inhibition, and (2) the curated KRAS core phosphoproteome described by Kabella et al. (*11*) after a 2-hr KRAS inhibitor treatment across three established PDAC cell lines. We observed strong enrichment of our KRAS^MUT^ upregulated phosphoproteome with the ERK-regulated phosphoproteome (*10*) at both 1-hr (enrichment=4.55, p=2.2×10^−30^) and 24-hr (enrichment=2.84, p=6.1×10^−34^) timepoints, as well as with the Kabella et al. (*11*) KRAS Core Signature (enrichment=6.44, p=1.46×10^−18^; **Fig. 4C**) (*10*). Notably, we observed strong overlap of upregulated (n=85) and downregulated (n=53) phosphoproteins compared to the Klomp et al. (*10*) 24-hr ERK-regulated dataset, likely reflecting greater convergence of the long-term perturbation with our steady-state conditions. Likewise, our KRAS^MUT^ downregulated phosphoproteome signature showed significant enrichment with the Klomp et al. (24-hr) dataset (enrichment=2.10, p=1.53×10^−8^) but no significant enrichment with the Klomp et al. (1-hr) dataset (p=7.64×10^−2^). Despite significant overlap in pairwise comparisons, across all datasets, ERBB2 (S1054) was the only phosphosite consistently downregulated, underscoring the narrow shared KRAS-dependent phosphoproteomic core signature across the studies (**Table S5**).

#### Comparative analysis between common PDAC KRAS mutations and other variants

Given that *KRAS^G12D^*, *KRAS^G12V^*, and *KRAS^G12R^* represent the most prevalent *KRAS* mutations in PDAC and have been associated with distinct biochemical properties (*4*) and clinical outcomes (*2, 20, 21*), we next sought to determine how their molecular profiles compare to those of less frequent alleles. We hypothesized that the molecular programs for KRAS^G12D^, KRAS^G12V^, and KRAS^G12R^ (DVR) would converge more strongly, suggesting an ideal cellular state that is compatible with PDAC development. We stratified our analysis by grouping DVR separately from the remaining KRAS variants (Other) and performed differential expression analysis relative to their isogenic KRAS^WT^ counterparts. Contrary to our hypothesis, we observed significant overlap between the upregulated and downregulated multi-omics signatures comparing DVR vs. Other (**Fig. 5A**). Functional enrichment analysis similarly showed a high degree of overlap between DVR and Other signatures across both the transcriptome and proteome, including suppression of interferon signaling responses (**Fig. 5B**). Furthermore, KMEA revealed upregulation of ERK1/2 and other MAPK family site motifs across both DVR and Other phosphoproteomic signatures (**Fig. 5C**). As before, DYRK kinase substrates were consistently downregulated in both groups, with this effect being more pronounced in the Other mutants compared to the DVR signature (**Fig. 5D**). In contrast, DVR downregulated sites – but not Other downregulated sites – were enriched for motifs corresponding to AMPK-related kinases downstream of the LKB1 tumor suppressor (BRSK, SIK, and MARK family kinases) (**Fig. 5D**). Together, these data reveal that transcriptomic, proteomic, and phosphoproteomic impacts largely converge irrespective of allele frequency in PDAC, arguing against a specific cell state induced by common KRAS variants in PDAC.

**Figure 5.**
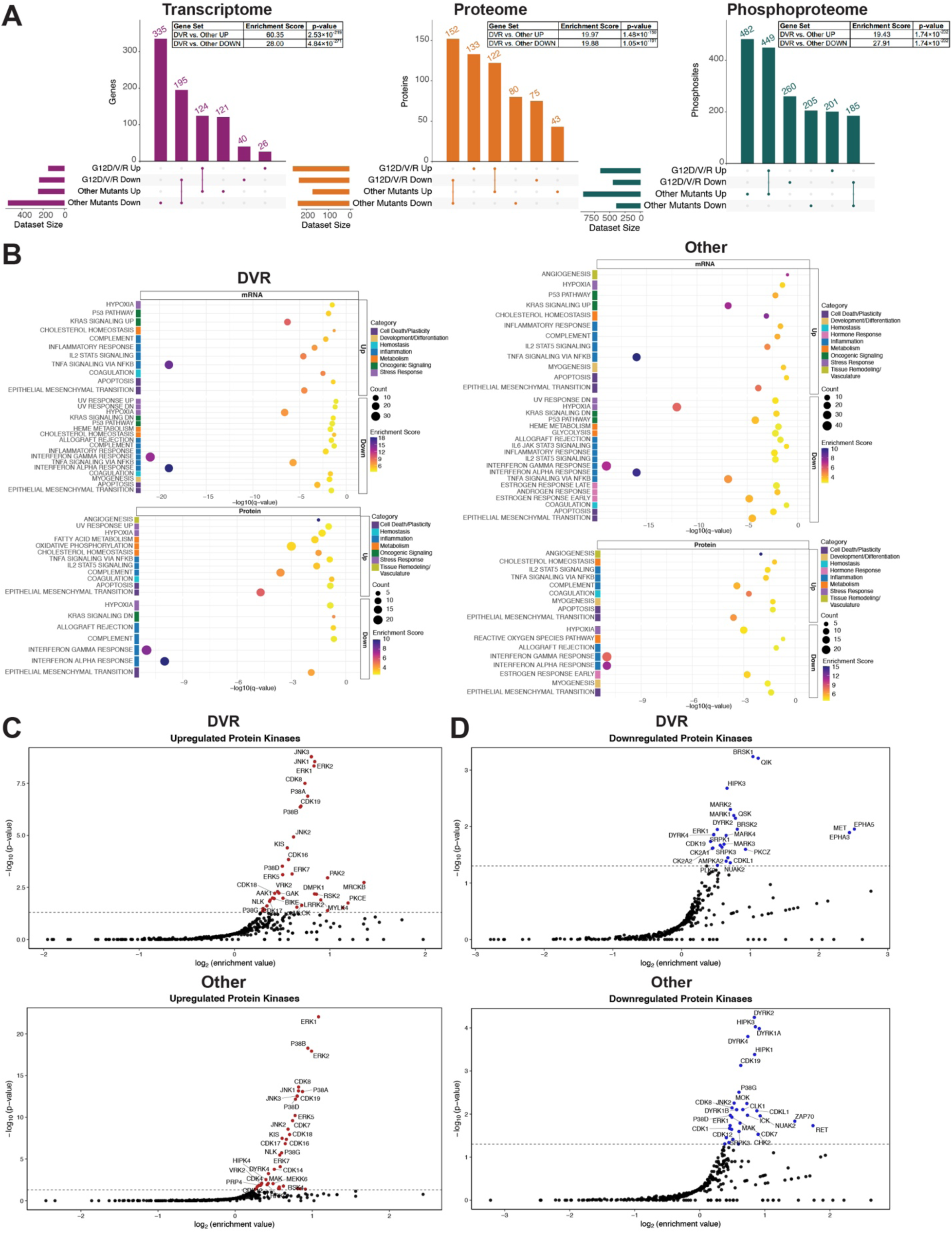
**Comparison of common versus less common KRAS variants in PDAC.** (A) UpSet plot illustrating overlap between the multi-omic signatures of DVR (KRAS^G12D^, KRAS^G12V^, and KRAS^G12R^) and Other (KRAS^G12C^, KRAS^G13C^, KRAS^Q61R^, and KRAS^Q61H^) KRAS variants. Enrichment scores and *p*-values of hypergeometric test are shown for corresponding upregulated (UP) or downregulated (DOWN) gene sets. (B) Pathway-level enrichment using the mSigDB Hallmarks dataset for the transcriptomic (mRNA) and proteomic (protein) differences (upregulated (Up) or downregulated (Down) for the DVR and Other multi-omic signatures each compared to KRAS^WT^. Color scale denotes enrichment score. Circle size designates gene count overlap. −log_10_(*q*-value) of hypergeometric test is plotted. (C) Kinase motif enrichment analysis (KMEA) using phosphosites in the DVR and Other multi-omic signatures reveals increased phosphorylation of ERK1/2 kinase substrates (upregulated protein kinases). Log_2_ enrichment vs. −log_10_(*p*-value) is plotted. (D) KMEA using phosphosites in the DVR and Other multi-omic signatures shows decreased phosphorylation if DYRK kinase substrates (downregulated protein kinases) for both signatures. AMPK-related kinases downstream of the LKB1 tumor suppressor (BRSK, SIK, and MARK family kinases) were uniquely downregulated in the DVR signature. Log_2_ enrichment vs. −log_10_(*s*-value) is plotted.

### Evaluation of KRAS mutant-selective signatures in PDAC

To uncover *KRAS* allele-specific biology in our datasets, we next focused on identifying mutant-selective signatures. After defining differentially expressed candidates relative to KRAS^WT^ for each KRAS mutant in each cell line, we classified them as mutant-selective if they were upregulated or downregulated in at least two of the four clones (score ≥ 2 or ≤ –2, respectively) for that mutant, while being differentially expressed in no more than one clone of all the other mutants (score ≤ 1 or ≥ –1). We visualized these candidate features (genes, proteins, or phosphosites) using heatmaps to assess whether allele-specific patterns emerged. Strikingly, the majority of allele-specific candidates showed similar directional change in candidate abundance with the other mutants such that no consistent mutant-specific transcriptomic, proteomic, or phosphoproteomic signatures stood out (**Fig. 6A**). Even for KRAS^Q61R^, which harbored the most mutant-selective candidates, the change in abundance for other mutants (relative to. KRAS^WT^) largely trended in the same direction (**Fig. 6B**), revealing that these candidate features (genes, proteins, phosphosites) were not exclusively enriched in KRAS^Q61R^-expressing cells. Consistent with this, supervised hierarchical clustering did not show specific clustering of Q61R across cell lines for KRAS^Q61R^ mutant-selective candidates (**Fig. 6C**), indicating that these candidate features are not truly KRAS^Q61R^-specific. Together, these findings suggest that cellular context (baseline gene expression and signaling networks), rather than intrinsic allele-specific properties, predominantly governs the molecular impact of KRAS mutations, thereby overriding subtle mutant-specific differences.

**Figure 6.**
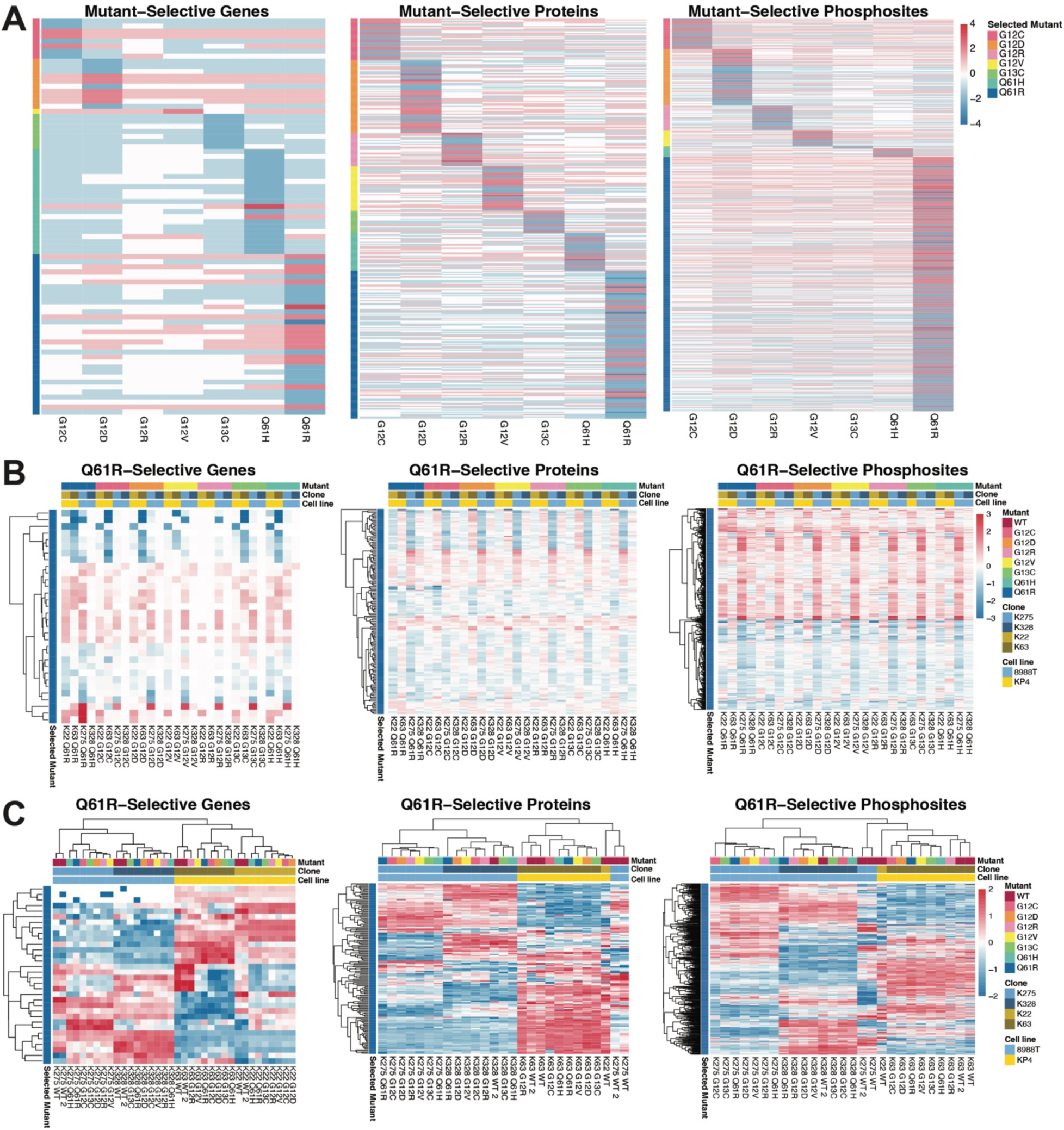
**Identification of mutant-selective signatures across multi-omic layers** (A) Heatmaps of transcriptomic, proteomic, and phosphoproteomic features (genes, proteins, phosphosites) meeting mutant-selective criteria (upregulated in ≥ 2 clones or downregulated in ≤ –2 clones for a given KRAS mutant and differentially expressed in ≤1 clone for other mutants (relative to WT) reveal limited allele specificity. Color scale is number of upregulated (+) or downregulated (-) clones (out of 4) for that feature (rows) of a given variant (columns). (B) Relative abundance (color scale denotes log_2_ fold-change relative to KRAS^WT^) for KRAS^Q61R^-selective candidates (in (**A**); rows) across all clones and mutant variants (columns) shows similarity in direction of expression across all layers. (C) Supervised hierarchical clustering analyses of KRAS^Q61R^-selective genes (transcriptome), proteins (proteome), and phosphosites (phosphoproteome) in (**A**). Color scale denotes row-normalized expression.

## DISCUSSION

In this study, we established an isogenic human PDAC cellular system to systematically interrogate the molecular consequences of seven different KRAS mutations on the transcriptome, proteome, and phosphoproteome. By restoring comparable KRAS expression in *KRAS*-deficient knockout clones, we created a platform that enabled direct allele-to-allele comparisons. Multi-omics profiling, while achieving excellent technical reproducibility, revealed that baseline cellular state, rather than intrinsic allele identity, was the predominant driver of variation in molecular signatures (**Fig. S5**). Overall, our datasets based on isogenic cells provide a high-quality multi-omic reference resource on the molecular impacts of common *KRAS* mutant variants in cancer.

A key strength of our approach is the ability to directly compare multiple KRAS alleles under controlled expression levels, minimizing confounding effects from variable KRAS abundance. However, a limitation of our system is that, by design, it relies on PDAC cell lines that have undergone signaling rewiring to survive in the absence of KRAS. It is therefore possible that KRAS plays a less dominant role in this context, contributing to the lack of global convergence across clones. Moreover, while our approach enables the simultaneous comparison of a broader panel of KRAS mutants, the limited number of available clones analyzed and the usage of *in vitro* cancer cell culture models may constrain our resolution to detect more subtle allele-specific effects. This interpretation aligns with prior reports of nuanced allele-specific differences, such as the inability of KRAS^G12R^ to regulate macropinocytosis (*7*). Nonetheless, macropinocytosis occurs even in KRAS^G12R^-expressing cells, arguing that compensatory, KRAS-independent pathways may sustain core cellular programs, thereby minimizing mutant-specific divergence.

To determine the generalizability of our method to define core KRAS-regulated pathways, we compared our KRAS^MUT^ signature with available KRAS reference signatures (*9–11, 17*). Our transcriptomic KRAS^MUT^ program showed modest overlap with the Hallmark KRAS gene set (*17*) and the KRAS-targeted siRNA-derived signatures of Klomp et al. (*9*), differences likely reflecting both the pan-cancer derivation of the Hallmark set and the acute perturbation design of the Klomp study when compared to our steady-state conditions in PDAC. Importantly, proteomic-level overlap was even more restricted, a finding that likely stems from biological factors – such as the imperfect correlation between mRNA and protein abundance – and differences in analytical sensitivity between RNA-Seq and DIA-MS platforms (*14, 15*). These results highlight that only limited KRAS-regulated core features are consistently maintained. Consistent with this, it has been postulated that specific KRAS mutants induce a “sweet spot” of signaling alterations to induce a cell state optimized for tumor development and maintenance in specific tissues (*22*). Indeed, the prevalence of *KRAS^G12R^* mutation exceeds the predicted frequency in PDAC (*23*), indicating that mutant-specific oncogenic signaling outputs may provide a selective advantage in a context-specific matter. However, contrary to this “goldilocks” hypothesis, common (DVR) and uncommon (Other) mutants largely converged on regulating the same signaling pathways and transcriptional outputs in our cells.

Pathway-level enrichment revealed that mutant KRAS expression suppressed interferon response genes at the transcriptome and proteome level, a finding that is consistent with previous reports of KRAS-mediated immune evasion in PDAC via suppression of type I interferon responses (*24*). At the phosphoproteome level, inferred ERK1/2 activity was consistently enriched across all KRAS mutants, whereas DYRK motif substrates were broadly suppressed. Our results align with a recent study that suggested that DYRK phosphosite motifs are upregulated as a compensatory response to sustained ERK inhibition in PDAC, CRC, and NSCLC models (*10*). Therefore, KRAS-induced MAPK signaling may suppress DYRK activity, however the precise mechanisms by which this occurs and whether DYRK signaling might enable bypass to KRAS inhibition remains to be elucidated.

Finally, attempts to identify mutant-specific signatures revealed that few transcripts, proteins, or phosphoproteins could be uniquely attributed to a single allele. These findings suggest that intrinsic allele-specific programs are modest in magnitude and are often overridden by baseline cell state. Although our analyses did not reveal robust allele-specific molecular signatures, prior clinical (*2, 20, 21*) and *in vivo* (*25–27*) studies have reported functional distinctions among KRAS mutants. However, recent evidence also suggests that these differences may be context-dependent rather than intrinsic. For instance, three-dimensional histological reconstruction and genetic mapping of low-grade pancreatic intraepithelial neoplasia (PanINs) – putative precursor lesions to PDAC – revealed that individual patients can harbor multiple *KRAS* hotspot mutations in spatially distinct PanINs and that more than one *KRAS* mutation can be detected within the same PanIN, indicating that different alleles can arise in parallel during early tumor development (*28*). Furthermore, the frequency of some mutations (*e.g.,* Q61) may be overrepresented in PanINs compared to PDAC (*28, 29*). These findings imply that extrinsic selective pressures such as cellular state, tissue context, and microenvironmental cues may amplify or constrain the subtle intrinsic biochemical differences between KRAS mutants throughout tumor evolution. Similarly, the emergence of a wide variety of secondary *KRAS* mutations in patients treated with KRAS inhibitors (*30–33*) suggests that diverse alleles can substitute for one another to mediate resistance. These data argue that the presence of mutant KRAS itself, rather than a specific allele, is sufficient to sustain oncogenic signaling of advanced cancer cells under therapeutic selection. Together, these observations argue that KRAS allele identity exerts a relatively minor influence on tumor cell behavior compared to the broader cellular and evolutionary context in which it operates. This implies that allele-specific phenotypes arise not from intrinsic KRAS differences alone, but from the interplay of these differences with selective pressures present in the tumor microenvironment that define cellular states. Ultimately, this work sets the stage for investigating how cellular state and microenvironmental cues intersect with KRAS allele identity to shape tumor biology and therapeutic response.

## MATERIALS AND METHODS

### Cell lines and cell culture conditions

Established human PDAC 8988T (ACC162) and KP4 (RCB1005) cell lines were obtained from Leibniz Institute DSMZ-German Collection of Microorganisms and Cell Cultures GmbH and RIKEN, respectively. 293FS* cells used for lentivirus production were a gift from Dr. N. Joshi. Ras-less mouse embryonic fibroblasts (MEFs) reconstituted with RAS and BRAF variants were acquired from the National Cancer Institute (NCI) RAS initiative. All cell lines were confirmed negative for mycoplasma using PCR testing by the Yale Molecular and Serological Diagnostics Core. Cell lines were maintained in DMEM (Corning, 10-013-CV) supplemented with 10% fetal bovine serum (FBS; Thermo Fisher Scientific, A5670701) and 1% penicillin/streptomycin (Thermo Fisher Scientific, 15140122).

### KRAS knockout clone generation

A single-guide RNA (sgRNA) targeting KRAS exon 2 (sequence: 5’-AATTACTACTTGCTTCCTGT-3’) was cloned into the CRISPR-Cas9 expression vector pSpCas9(BB)-2A-GFP (PX458, Addgene plasmid #48138). 1×10^6^ 8988T and KP4 cells were nucleofected (Amaxa) with the sgKRAS-Cas9-2A-GFP construct. Forty-eight hours later, GFP-positive cells were isolated by flow cytometry using a BD FACSymphony™ S6 Cell Sorter (BD Biosciences) to enrich for vector-expressing populations. After seven days, GFP-negative cells were single-cell sorted to generate stable 8988T and KP4 clones, yielding approximately 680 single-cell isolates for each parental cell line. Approximately 70 KP4 clones and 100 8988T clones were initially screened for KRAS expression by immunoblotting. Of the 4 KP4 and 40 8988T clones clearly lacking detectable KRAS protein, all KP4 clones and 13 8988T clones were further validated by PCR amplification of KRAS exon 2, as previously described, and amplicon next-generation sequencing using the Massachusetts General Hospital (MGH) CCIB DNA Core. 3 of 4 candidate KP4 clones and 12 of 13 candidate 8988T clones were confirmed as *KRAS* knockouts (KO), each harboring out-of-frame indels at the sgRNA target site. From these, KO22 and KO63 (KP4) and KO275 and KO328 (8988T) were selected for downstream experiments.

### Genomic DNA isolation and sequencing

Genomic DNA was extracted from *KRAS* KO clones and the parental cell line using the High Pure PCR Template Prep Kit (Roche, 11796828001). Whole-exome sequencing libraries were prepared by the Yale Center for Genome Analysis (YCGA) using the IDT xGen V2 Exome Enrichment Kit and sequenced on the Illumina HiSeq platform (paired-end, 100-bp reads) to a mean target coverage of approximately 30×.

### Generation of isogenic KRAS variant-expressing PDAC cell lines

To make KRAS expression constructs (LV-PGK-eGFP-2A-KRAS), lentiviral backbone vector (LV 1-5, Addgene #68411) was linearized by restriction enzyme digestion at designated sites. The PGK promoter and GFP cassette were PCR-amplified from the MSCV Puro-pGK:GFP construct (Addgene #68486) and inserted into the linearized lentiviral backbone using Gibson Assembly (New England Biolabs (NEB), #E2611) as previously described (*34*). KRAS mutant cDNA sequences were PCR-amplified from the RAS Mutant Clone Collection (Addgene, Kit #1000000089) with Q5 Hot Start High-Fidelity DNA Polymerase (NEB, M0494S) using forward primers containing a NheI-HF (NEB, R3131S) or BamHI-HF (NEB, R3136S) restriction digest site and a P2A self-cleaving peptide and the reverse primer containing an AscI (NEB, R0558S) restriction digest site. PCR products were gel-purified (Qiagen, 28706), digested with respective restriction enzymes, PCR purified (Qiagen, 28106), and ligated with the digested backbone for 1-2 hours using T4 ligase (NEB, M0202S). The *KRAS^Q61H^* insert was generated via overlapping PCR to introduce the desired point mutation, followed by restriction digestion and cloning into the lentiviral backbone using the same restriction enzyme and ligation strategy described for the other *KRAS* variants. The ligation products were transformed into Stbl3 competent *E. coli* (Invitrogen, C737303) and selected on ampicillin-containing agar plates. Individual positive clones were cultured overnight in LB Broth (Thermo Fisher Scientific, 12780052). Plasmids were purified using a QIAprep Miniprep Kit (Qiagen, 27106) and validated by Sanger sequencing performed by the W. M. Keck Biotechnology Resource Laboratory at Yale. To produce lentivirus, lentiviral backbone, packaging vector (psPAX2, Addgene #12260), and envelope vector (VSV-G, Addgene #8454) were co-transfected into 293FS* cells using TransIT-LT1 (Mirus Bio, MIR2306). Culture media was changed 16 hours post-transfection. Supernatants were collected at 48- and 72-hours post-transfection, filtered through 0.45 µm syringe filters, and used immediately or store at −80 °C prior to use. For stable KRAS expression, KRAS-deficient clones were seeded ≥16 hours prior to transduction and treated with lentiviral supernatant (LV-PGK-eGFP-2A-KRAS) supplemented with 8 μg/mL polybrene (EMD Millipore, TR-1003-G). KRAS-reconstituted cells were sorted based on fluorophore intensity using a FACSymphony™ S6 Cell Sorter, wherein GFP fluorescent intensity was used as a proxy for desired KRAS expression. Cells were replated after sorting and stable fluorescence was confirmed 7 days post sort. Expression of KRAS protein was confirmed by immunoblotting.

### Immunoblotting

7.5×10^5^ cells were seeded per well in 6-well plates the day prior to collection of whole cell protein lysates. For collection, cells were rinsed with cold 1X PBS, scraped, pelleted, and resuspended in protein lysis buffer consisting of Pierce® RIPA buffer (Thermo Fisher Scientific, 89900) supplemented with 1:100 0.5M EDTA (Thermo Fisher Scientific) and 1:100 Halt™ protease/phosphatase inhibitor cocktails (Thermo Fisher Scientific; 78444), and incubated for 15 minutes at 4 °C in a rotator. Cells were then pelleted at maximum speed (∼14,000 × g) for 15 minutes at 4 °C, and supernatant containing the protein lysate was collected and transferred to a new microcentrifuge tube. Protein concentrations were quantified using the Pierce® BCA Protein Assay Kit (Thermo Fisher Scientific, 23225) following the manufacturer’s instructions. Equal amounts of protein (40 μg) were mixed 1:1 (by volume) with 2X Laemmli Sample Buffer (Bio-Rad, 1610737) containing 5% β-mercaptoethanol (Thermo Fisher Scientific, 21985023), boiled at 95 °C for 5 minutes, loaded onto Mini-PROTEAN® 4–20% TGX™ stain-free precast gels (Bio-Rad), and separated by SDS-PAGE. Proteins were transferred onto nitrocellulose membranes using the Trans-Blot® Turbo™ Transfer System (Bio-Rad). Membranes were blocked for 1 hour at room temperature in Intercept® (PBS) Blocking Buffer (LI-COR, 927-70001) diluted 1:2 in phosphate-buffered saline (PBS) or 5% Omniblok non-fat dry milk powder (AmericanBio, AB10109-01000) in Tris-buffered saline (TBS). Membranes were then incubated overnight at 4 °C with primary antibodies diluted in either Intercept Blocking Buffer with 0.1% Tween-20 (PBS-T, 1:2) or in TBS-T with 5% non-fat dry milk powder. The following antibodies were used for immunoblotting: rabbit anti-HSP90 (Cell Signaling Technologies (CST), 4877, 1:10,000), rabbit anti-pERK1/2 (T202/Y204) (CST, 4370, 1:2000), mouse anti-ERK1/2 (CST, 9107, 1:1000), rabbit anti-pAKT (S473) (CST, 4060, 1:1000), mouse anti-AKT (CST, 2966, 1:2000), mouse anti-KRAS (Sigma-Aldrich, 3B10-2F2, 1:1000). Primary rabbit antibodies were detected with antirabbit IgG (H+L) (DyLight™ 800 4X PEG Conjugate) (CST, 5151, 1:10,000). Primary mouse antibodies were detected with anti-rabbit IgG (H+L) (DyLight™ 800 4X PEG Conjugate) (CST, 5151, 1:10,000), anti-mouse IgG (H+L) (DyLight™ 680) (CST, 5470, 1:10,000), or anti-mouse IgGκ light chain-horseradish peroxidase (HRP) (Santa Cruz Biotechnology, sc-516102; 1:1,000). Chemiluminescent detection was performed using the SuperSignal™ West Dura Extended Duration Substrate (Thermo Fisher Scientific, 34076X4). Fluorescence or chemiluminescence images were acquired under non-saturating exposure conditions using the ChemiDoc imaging system (Bio-Rad). Precision Plus Protein Kaleidoscope Prestained Protein Standards (Bio-Rad, 1610375) were used as protein size markers for individual blots. The resulting images were saved in Image Lab software (v6.1.0; Bio-Rad) format for subsequent densitometric quantification.

### RNA isolation and sequencing

7.5×10^5^ cells were seeded per well in 6-well plates the day prior to collection. RNA was extracted using the Qiagen RNeasy Mini kit (Qiagen, 74106) following the manufacturer’s protocol with RNase-free DNAse I treatment (Qiagen, 79254) to remove any genomic DNA contamination. Only high-quality RNA samples (RIN>7), confirmed using an Agilent Bioanalyzer, were used in RNA-seq analyses. Bulk RNA-seq libraries were prepared via polyA selection (Illumina) and sequenced (>25M 100-bp paired-end (2×100) reads on NovaSeq (Illumina) through the YCGA.

### Protein extraction and digestion for mass spectrometry

For proteomics and phosphoproteomics analyses, 10M urea proteomics/phosphoproteomics lysis buffer was prepared by dissolving 10M Urea (Thermo Fisher Scientific, 29700), 100 mM ammonium bicarbonate BioUltra (Sigma Aldrich, 1066-33-7), and Complete™ Protease Inhibitor Cocktail (Sigma-Aldrich, 11873580001) in HPLC-grade water (Fisher Chemical, 7732-18-5) and stored at −80 °C. Approximately 4×10^6^ cells were plated in triplicate in 10-cm dishes the day prior to protein lysate collection. Cells were rinsed twice with cold 1X PBS, scraped, and pellets were reconstituted in 250 uL of 10M urea proteomics/phosphoproteomics lysis buffer containing freshly added 1:100 Halt Phosphatase Inhibitor Cocktail (Thermo Fisher Scientific, 78420). Cell lysates were sonicated twice for 1 min at 4 °C using a VialTweeter device (Hielscher-Ultrasound Technology). The lysates were then centrifuged at 20,000 × g for 1 hour to pellet insoluble material. Supernatant containing the protein lysate was collected and transferred to a 2 mL microcentrifuge tube. Protein concentration was quantified with the Bio-Rad protein assay. Proteins were reduced with 10 mM DTT at 56 °C for 1 hour, followed by alkylation with 20 mM iodoacetamide (IAA) in the dark at room temperature for 1 hour. The reduced and alkylated proteins were then digested using a precipitation-based protocol, as previously described (*35*). Briefly, proteins were mixed with five volumes of cold precipitation solution (50% acetone, 50% ethanol, 0.1% acetic acid) and incubated at −20 °C overnight. Precipitated proteins were pelleted by centrifugation at 20,000 × g for 40 min, washed with precipitation solution, and centrifuged again under the same conditions. After removing the precipitation solution, residual solvent was evaporated in a SpeedVac (Thermo Fisher Scientific). Proteins were then digested overnight at 37 °C with sequencing-grade porcine trypsin (Promega, V5111; 1:50 enzyme-to-substrate ratio) in 300 μl of 100 mM ammonium bicarbonate. The peptide mixture was desalted on C18 MacroSpin columns (NEST Group, Inc.) following the manufacturer’s protocol and quantified using a NanoDrop spectrophotometer (Thermo Fisher Scientific). Two dish replicates were processed for each condition resulting in 80 samples for the total proteome and phosphoproteome analysis.

### Phosphopeptide enrichment

Phosphopeptides were enriched using the High-Select™ Fe-NTA kit (ThermoFisher Scientific, A32992) following the manufacturer’s protocol (*36*). Briefly, the peptide–resin suspension was incubated for 30 min at room temperature with gentle shaking every 10 mins, then transferred to a filter tip (TF-20-L-R-S, Axygen) and centrifuged to discard the flow-through. The resin was then washed three times with 200 µL wash buffer (80% acetonitrile, 0.1% trifluoroacetic acid) and twice with 200 µL H₂O. Phosphopeptides were eluted twice with 100 µL elution buffer (50% acetonitrile, 50% NH₃·H₂O), dried in a SpeedVac, and quantified by NanoDrop.

### Mass spectrometry measurements

For liquid chromatography-mass spectrometry (LC-MS) analysis, peptides were analyzed as described previously (*35, 37*). Peptide separation was carried out using an EASY-nLC 1200 system (ThermoFisher Scientific) with a self-packed PicoFrit column (New Objective; 75 μm × 50 cm) containing ReproSil-Pur 120A C18-Q 1.9 μm resin (Dr. Maisch GmbH, Ammerbuch, Germany). Peptides were eluted over a 150-min gradient from 4% to 37% buffer B (80% acetonitrile, 0.1% formic acid) with buffer A (0.1% formic acid in water) at a flow rate of 300 nl/min, while maintaining the column at 60 °C (PRSO-V1 oven, Sonation GmbH). Eluted peptides were analyzed on an Orbitrap Fusion Lumos Tribrid mass spectrometer (ThermoFisher Scientific) equipped with a NanoFlex ion source (spray voltage 2000 V, capillary temperature 275 °C). The data independent acquisition mass spectrometry (DIA-MS) acquisition consisted of an MS1 survey scan followed by 33 variable-window MS2 scans, as described previously (*38, 39*). MS1 scans were acquired from 350–1650 m/z at a resolution of 120,000 (m/z 200) with an AGC target of 2.0E6 and a maximum injection time of 50 ms. MS2 scans were collected at a resolution of 30,000 (m/z 200) using HCD with 28% normalized collision energy, an AGC target of 1.5E6, and a maximum injection time of 52 ms. The default precursor charge state was set to 2, and both MS1 and MS2 spectra were recorded in profile mode.

### Quantification and Statistical Analysis

#### Immunoblot quantification and statistical analysis

Protein quantification from immunoblots was performed using Image Lab software (v6.1.0; Bio-Rad). Bands corresponding to the protein of interest were defined using the Lane and Band detection tools, and background subtraction was applied using the local background method. Integrated adjusted band intensities were initially normalized to the corresponding loading control (HSP90). Phosphorylated AKT (pAKT) and ERK1/2 (pERK1/2) were then normalized to their non-phosphorylated counterparts AKT and ERK1/2, respectively. All samples were then normalized to KRAS^WT.^ Normalized ratios from n = 3 biological replicates are presented as mean ± SEM (standard error of the mean) or geometric mean ± geometric SD (standard deviation) in bar graphs and then analyzed using one-way repeated measures ANOVA with Tukey’s or Dunnett’s post-hoc test or Brown-Forsythe lognormal ANOVA with Games-Howell’s post-hoc test in Prism v10.6.0 (Graphpad Software). *p* < 0 .05 was used as level of significance for all statistical analyses.

#### Whole exome sequencing analysis and variant detection

Raw whole exome sequencing data were processed on the Galaxy web platform to generate analysis-ready BAM files. Preprocessing steps included read filtering using Filter BAM datasets on a variety of attributes (Galaxy v2.4.1), duplicate removal using RmDup (Galaxy v2.0.1), indel left-alignment using BamLeftAlign (Galaxy v1.3.1), and recalculation of MD tags and mapping qualities using CalMD (Galaxy v2.0.2). The resulting CalMD-corrected BAM files from each CRISPR-edited clone were paired with the corresponding parental cell line BAMs, which served as matched normal controls, for somatic variant calling against the hg19 reference genome.

Somatic variant detection was performed using the Galaxy Somatic Variant Calling workflow incorporating VarScan Somatic (Galaxy tool v2.4.3.6). VarScan was executed with customized variant-calling parameters, including a minimum base quality of 28 and a minimum mapping quality of 1, while all other calling and posterior filtering parameters were retained at their default settings. Normal sample purity and tumor sample purity were both explicitly specified as 1.0, reflecting the clonal nature of the cell line samples. Under VarScan’s somatic classification criteria, variants were considered somatic if the tumor variant allele frequency (VAF) was ≥10%, the normal VAF was ≤2%, minimum depth and supporting read thresholds were met, and the somatic *p*-value was ≤0.05. Identified single-nucleotide variants (SNVs) and indels were functionally annotated using SnpEff (Galaxy v4.3+T.galaxy1) and further annotated with information from ClinVar, dbSNP, the COSMIC Cancer Gene Census (CGC), and the Cancer Genome Interpreter (CGI). Coding variants were summarized in a protein-effect table reporting allele frequencies, predicted amino-acid substitutions, and cancer-associated gene annotations. The Galaxy workflow was adapted from the Galaxy Training Network tutorial “Identification of somatic and germline variants from tumor and normal sample pairs” (https://training.galaxyproject.org/training-material/topics/variant-analysis/tutorials/somatic-variants/tutorial.html).

Copy-number variation (CNV) analysis was performed in R (v4.3.2) using the Bioconductor package cn.mops (v1.48.0), with supporting packages GenomicRanges (v1.54.1) and GenomicFeatures (v1.54.4). Read depth was quantified across hybrid-capture target regions extended by ±30 bp, and an exome-optimized model was used to infer copy-number gains and losses. CNV segments were overlapped with gene coordinates from the UCSC hg19 reference annotation, and affected genes were cross-referenced with curated cancer gene resources including COSMIC CGC, ClinVar, and CGI.

#### RNA-sequencing analysis

Low-quality bases and adapter sequences were trimmed using Trim Galore (v0.6.10). Cleaned reads were aligned to the Gencode human genome (Release 48; version GRCh38.p14) using the STAR aligner (v2.7.7a). Differential gene expression analysis was performed using DESeq2 (v1.46.0) (*40*). Data was prefiltered to remove low-expressing genes (≤10). Genes with a fold change of ≥1.3 and an adjusted *p*-value ≤ 0.05 were considered differentially expressed. Three comparisons were performed separately for each clone: (i) each KRAS mutant (n = 1) was tested against two KRAS wild-type (KRAS^WT^) controls (n = 2) to find mutant-specific signatures, (ii) all samples from the KRAS mutant conditions (n = 7) were tested against KRAS wild-type controls (n = 2) to find pan-mutant signatures, and (iii) all samples from the “DVR” subtype (KRAS^G12D^, KRAS^G12V^, and KRAS^G12R^; n = 3) and “Other” subtype (KRAS^G12C^, KRAS^G13C^, KRAS^Q61H^, KRAS^Q61R^; n = 4) were tested against KRAS^WT^ controls (n = 2) to find DVR signatures. All RNA-seq analyses were conducted in R (v4.4.2).

#### Mass spectrometry data processing

Label-free data were analyzed in Spectronaut v19 using the directDIA+ (*39*) workflow against the human SwissProt database supplemented with manually added KRAS mutant sequences (20,426 entries, downloaded March 2025), applying default Spectronaut settings (*15*). For total proteome searches, carbamidomethylation of cysteine was set as a fixed modification, while methionine oxidation and N-terminal acetylation were specified as variable modifications. In phosphoproteome searches, phosphorylation of serine, threonine, and tyrosine (S/T/Y) was additionally included as a variable modification. Peptide- and protein-level FDRs were controlled at 1%, and data were filtered by Qvalue. PTM site localization in Spectronaut v14 was performed using the PTM function with a score cutoff of >0.75 (*41, 42*). All other Spectronaut parameters were left at default, with “Interference Correction” enabled and “Global Normalization (Median)” applied. The precursor and experiment-wide protein Q-values were set to 0.01, while the run-wise protein Q-value was 0.05. Protein and peptide quantification were based on the top 3 precursors and top 3 stripped peptide sequences, respectively. Total proteome data were exported via the protein pivot report, using PG.ProteinGroups as the unique identifier and PG.Quantity for quantification. Phosphoproteome data were exported via the peptide pivot report, with EG.PrecursorId as the unique identifier and EG.TotalQuantity as the quantification column. Relative intensities below 10 were replaced with NA, followed by log₂ transformation and normalization using the limma::normalize.cyclic.loess() function (*43*). For phosphoproteome analysis, precursor-level pivot reports were exported from Spectronaut using two PTM localization score cutoffs (0.75 and 0), following the strategy outlined in our previous work (*15*). Briefly, the first report (localization probability > 0.75; class I sites (*13*); n = 116,086 unique modified precursors) was used to identify confidently localized phosphopeptides. The second report (localization probability > 0; n = 181,427 unique precursors) was restricted to the precursors from the first report and served to extract intensity values. To define representative phosphopeptides (phos.id), we selected: (i) the precursor with the greatest number of valid values across the 80 MS runs, and (ii) if tied, the precursor with the highest summed intensity. This yielded 63,448 unique phos.ids from 5,967 phosphoprotein groups, preserving information on multiply phosphorylated peptides for downstream analyses. These mapped to 56,768 unique phosphorylation sites. Phosphoprecursor intensities below 8 were excluded, and the data were log₂ transformed and normalized using the limma::normalize.cyclic.loess() function (*43*). Two samples with low quality (number of quantified unique phos.ids below 20,000; both replicates of K22_G13C) were repeated in another batch replicate, leading to an improved and comparable coverage (∼37,000 IDs). However, even after a batch effect removal attempt using harmonizR (*44*), the repeated sample showed a notable experimental batch effect and thus was rigorously reported but excluded from downstream analysis. Sequence windows flanking the modification site (15-mers) were extracted using the PTMoreR package (1.1.0) (*45*). To confirm comparable KRAS expression across variants using our DIA-MS data, we extracted precursor intensities for KRAS peptides and filtered them to exclude sequences mapping to NRAS or HRAS. Seven unique KRAS-specific peptide precursors were identified and a representative peptide consistently detected across all samples was used to confirm comparable KRAS abundance across variants.

#### Statistical analysis of proteomics data

Statistical analysis was performed in R using the limma package (3.58.1) (*46*) with the standard workflow (lmFit(), contrasts.fit(), eBayes()). Multiple testing correction was applied using the Benjamini–Hochberg FDR method, with significance defined as FDR < 0.05 and absolute fold change > 1.3. Moderated log₂ fold changes from limma were used for downstream analyses. Three statistical analyses were performed separately for each clone: (i) each of the KRAS^MUT^ condition (n = 2) was tested against two KRAS^WT^ controls (n = 4); data were prefiltered to contain at least 2 valid values in each condition tested; this analysis was used to find mutant-specific signatures, (ii) all samples from the KRAS^MUT^ conditions (n = 14) were tested against KRAS^WT^ controls (n = 4); data were prefiltered to contain at least 3 valid values in each condition tested; this analysis was used to find pan-mutant signatures, and (iii) all samples from the “DVR” subtype (KRAS^G12D^, KRAS^G12V^, and KRAS^G12R^; n = 6) and “Other” subtype (KRAS^G12C^, KRAS^G13C^, KRAS^Q61H^, KRAS^Q61R^; n = 8) were tested against KRAS^WT^ controls (n = 4); data were prefiltered to contain at least 3 valid values in each condition tested; this analysis was used to find DVR signatures.

#### Identification of pan-mutant and mutant-selective KRAS signatures

Following differential expression analysis, each gene, protein, or phosphosite was assigned a score of +1 or –1 per cell line depending on whether it met the statistical significance threshold and the direction of regulation. Scores were then summed across cell lines to generate an overall score for each feature, which was subsequently used to define KRAS signatures. For the pan-mutant KRAS, DVR, and Other signatures, candidate hits were defined as upregulated (overall score ≥ 2) or downregulated (overall score ≤ –2) in at least two of the four cell lines. To identify mutant-selective signatures, a gene, protein, or phosphosite was determined as specific to a given KRAS mutant if it (1) was consistently differentially expressed in at least two clones of that allele, with the sign of the score reflecting directionality, and (2) was present in no more than one clone of the other alleles.

#### Enrichment analysis

A multiple gene list enrichment analysis was performed using the Metascape web interface (https://metascape.org) (*47*) to perform functional enrichment analysis against a custom background gene set consisting of unique genes/proteins/phosphoproteins included in the corresponding statistical analysis. Over-representation analysis of selected signature gene sets was performed using the phyper() function in R. The following sets were used: (i) the KRAS-dependence gene set based on Klomp et al. (*9*), (ii) the KRAS Hallmarks genes set based on the MSigDB (*17*), (iii) the 1-hr and 24-hr ERK-dependent KRAS phosphoproteome based on Klomp et al. (*10*), and (iv) the core KRAS signaling signature based on Kabella et al. (*11*).

#### Kinase motif enrichment analysis

Phosphoproteomics datasets were filtered to include only peptides with a single site of phosphorylation. Enriched motifs in sites significantly upregulated or downregulated in at least two clones within the pan-KRAS^MUT^ group, “DVR” KRAS group, or two “Other” KRAS mutant group compared to matched KRAS^WT^-expressing cells were identified using the phosphoproteomic enrichment analysis tool (kinase-library.mit.edu) with predetermined foreground and background sets. Sites were called as conforming to a motif if their percentile rank was within the top 15 Ser-Thr kinases or the top 8 Tyr kinases, and enrichment *p*-values were adjusted by Benjamini-Hochberg procedure.

#### Data visualization

Hierarchical clustering analysis and heatmap visualization was performed using the R package pheatmap (1.0.13). UpSet plots were generated using the R package UpSetR (1.4.0). Correlation plots were generated using the R package corrplot (0.95). Barplots were generated using the R package ggplot2 (3.5.2).

## ACKNOWLEDGEMENTS

We thank the Muzumdar and Liu lab members for helpful discussions and invaluable feedback; X. Ge, J. Singh., S. S. Agabiti, and W. Li for technical support; C. F. Ruiz for assistance with RNA-seq analysis; the Yale Center for Genome Analysis (YCGA) for library preparation, RNA-sequencing, and whole exome sequencing; the W. M. Keck DNA Sequencing Facility at Yale and MGH CCIB DNA Core for DNA sequencing; F. Fenteany and the Yale West Campus Flow Cytometry Facility for flow cytometry assistance; N. Joshi, the National Cancer Institute (NCI) RAS Initiative, DSMZ, and RIKEN for cell lines; and T. Jacks, F. Zhang, and the NCI Ras Initiative for plasmid constructs.

## Author Contributions

Y.Q.A and M.D.M. conceived of the project and designed the study. Y.Q.A. designed and performed studies in cell lines including molecular cloning, establishing isogenic cell lines, data acquisition, RNA-sequencing data analysis, and interpretation. B.S. analyzed MS data and conducted quantitative and statistical analysis of proteomics and phosphoproteomics data, and enrichment analysis for all datasets. C.S.M. established the *KRAS* knockout cell lines, performed whole exome sequencing analysis, and provided cloning assistance. Y.D. carried out mass spectrometry sample preparation and performed LC-MS/MS data acquisition. B.E.T. performed kinase substrate enrichment analyses. Y.S.L. and M.D.M. supervised the overall study. Y.Q.A. and M.D.M wrote the manuscript with input from all authors.

## Data and Materials Availability

The mass spectrometry data and raw output tables have been deposited in the ProteomeXchange Consortium via the PRIDE (*48*) partner repository with the dataset identifier PXD071086. RNA-sequencing data have been deposited in NCBI’s Gene Expression Omnibus and are accessible through GEO Series accession number GSE314073. Plasmids generated in this study will be deposited in Addgene. All other unique reagents are available upon request.

## SUPPLEMETARY FIGURES

**Figure S1.**
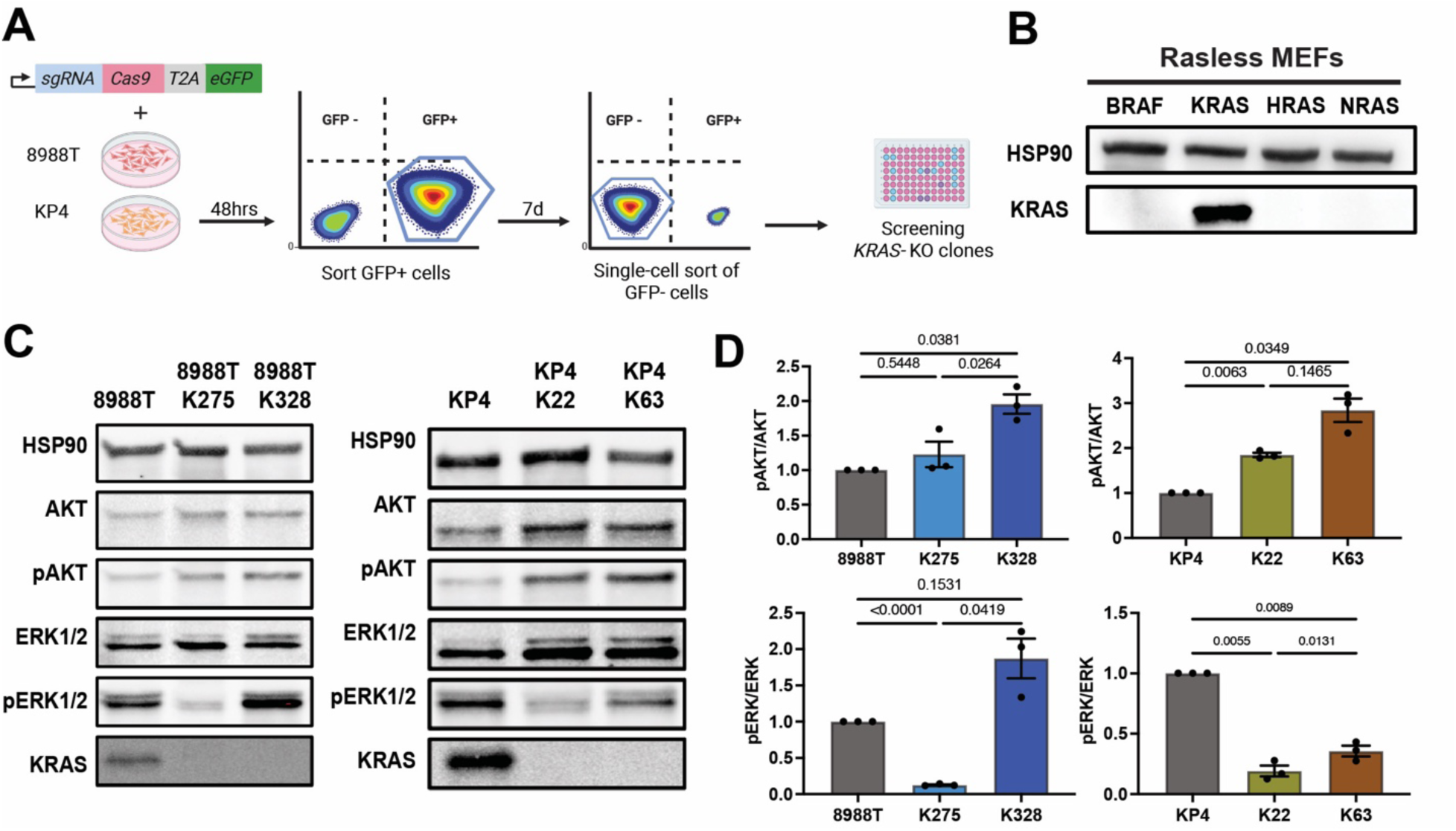
**Generation of *KRAS* knockout clones.** (A) Schematic of the CRISPR/Cas9 strategy used to generate *KRAS* knockout 8988T and KP4 cell lines by targeting exon 2 of *KRAS*. (B) Immunoblot of RAS*-*less mouse embryonic fibroblasts (MEFs) reconstituted with BRAF^V600E^, KRAS^WT^, HRAS^WT^, or NRAS^WT^, confirming specificity of the anti-KRAS antibody (clone 3B10-2F2). (C) Immunoblots showing absence of KRAS protein in 8988T (K275, K328) and KP4 (K22, K63) *KRAS* knockout clones and baseline PI3K (pAKT) and MAPK (pERK1/2) levels relative to parental cell lines. Loading control is HSP90. Images are representative of n = 3 biological replicates. (D) Bar graphs show quantified pERK/ERK and pAKT/AKT levels relative to the parental cell line (8988T or KP4) (mean ± SEM of n = 3 biological replicates) of immunoblots in (**C**). *p*-values of repeated measures one-way ANOVA with Tukey’s post hoc test are shown.

**Figure S2.**
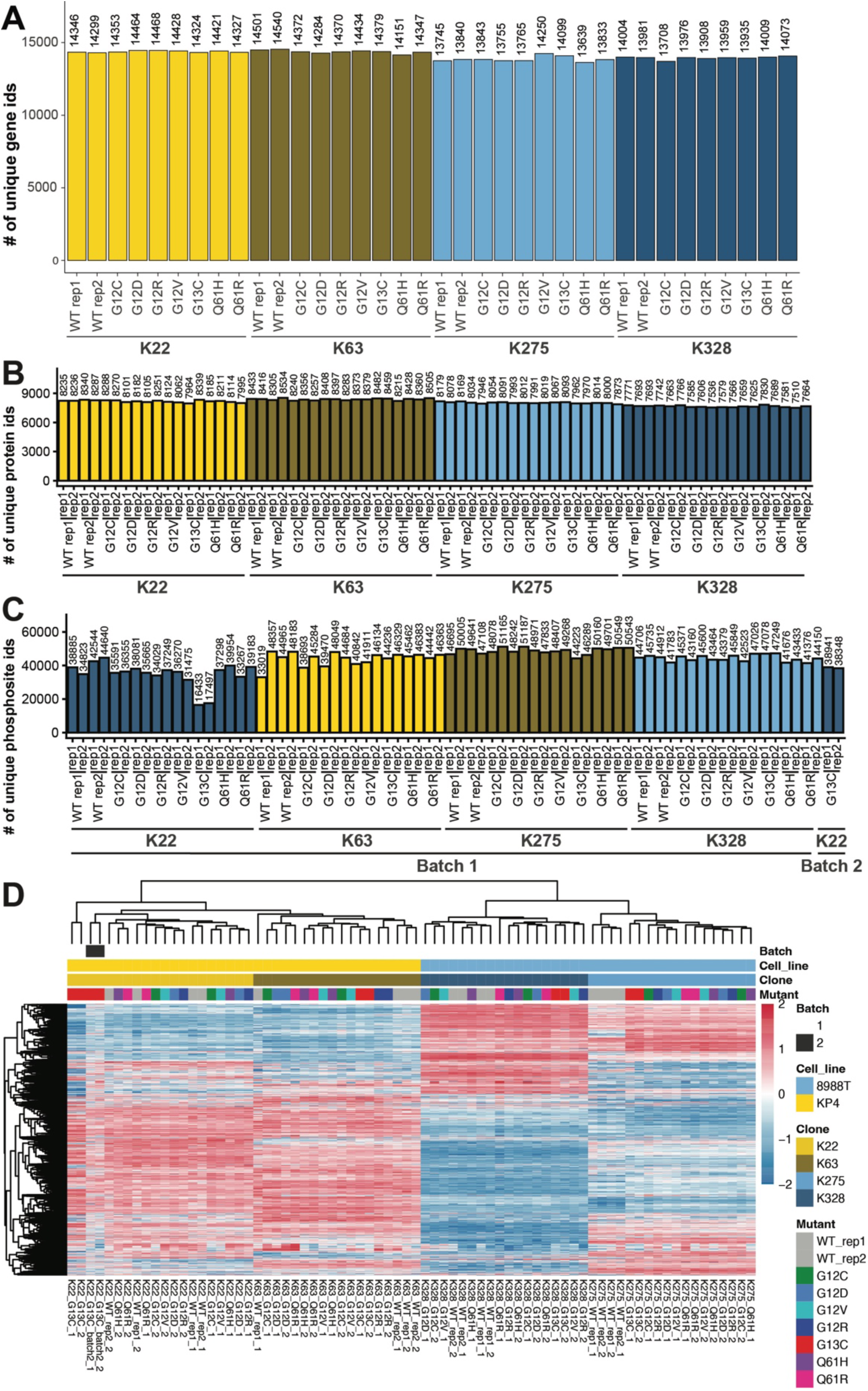
**Depth of omics coverage across molecular layers.** (A) Total number of protein-coding genes detected across all samples analyzed via RNA-seq. (B) Total number of proteins detected across all samples analyzed via DIA-MS. Data from each of two technical (dish) replicates are shown. (C) Total number of phosphopeptides detected across all samples analyzed via DIA-MS. Data from each of two technical (dish) replicates is shown. **K22 KRAS^G13C^ was re-acquired separately in a second batch (Batch 2).** Data from each of two technical (dish) replicates are shown. (D) Hierarchical clustering analysis (HCA) of complete phosphosite cases following batch-effect correction of additional K22 KRAS^G13C^ replicates (Batch 2). Both Batch 2 samples cluster away from all other high-quality Batch 1 K22 samples, consistent with persistent batch effects following data integration. As a result, re-acquired samples were excluded from downstream analyses. Data from each of two technical (dish) replicates are shown.

**Figure S3.**
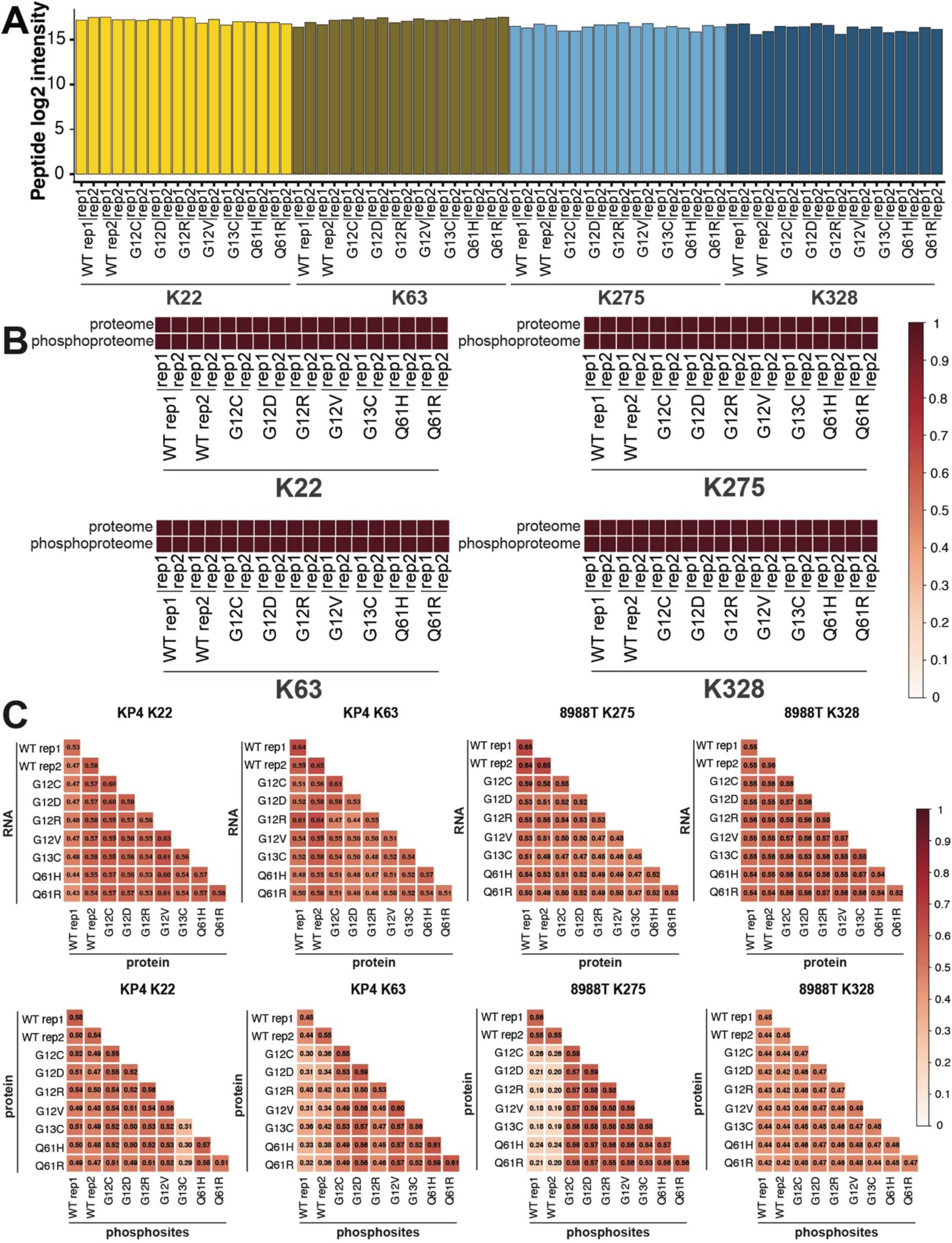
**Reproducibility and cross-layer concordance of multi-omics measurements.** (A) Quantification of a representative KRAS-specific peptide precursor (_VKDSEDVPMVLVGNK_.2) consistently detected across all samples confirms similar KRAS expression across variants. Peptides were filtered to exclude those mapping to NRAS or HRAS. (B) Pairwise Spearman absolute correlation analysis of technical replicates demonstrates high reproducibility of both the proteome and phosphoproteome datasets. Color scale represents Spearman’s ρ (rho) values. (C) Pairwise Spearman correlations between mRNA and protein abundance and between protein and phosphosite relative to the average of all samples. Color scale represents Spearman’s ρ (rho) values.

**Figure S4.**
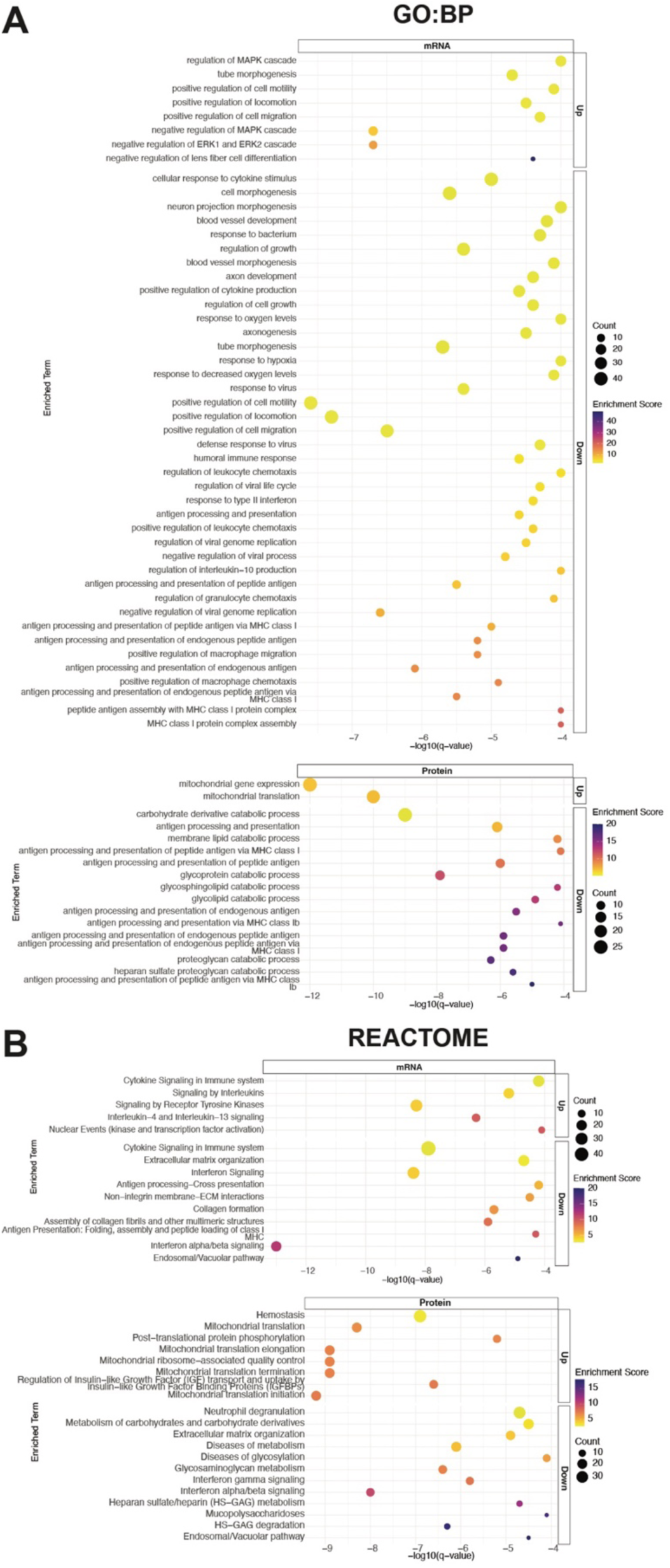
**Over-representation analysis of enriched pathways across transcriptomic and proteomic layers.** (A) Pathway-level enrichment using the Gene Ontology Biological Process (GO: BP) for the transcriptomic (mRNA) and proteomic (protein) differences (upregulated (Up) or downregulated (Down)) for the KRAS^MUT^ multi-omic signatures defined by selective differentially expressed hits (FC ≥ 1.3 and FDR ≤ 0.05) in at least 2 out of 4 clones (≥2 clones) compared to KRAS^WT^. Color scale denotes enrichment score. Circle size designates gene count overlap. −log_10_(*q*-value) of hypergeometric test is plotted. (B) Pathway-level enrichment using the Reactome pathways for the transcriptomic (mRNA) and proteomic (protein) differences (upregulated (Up) or downregulated (Down)) for the KRAS^MUT^ multi-omic signatures

**Figure S5.**
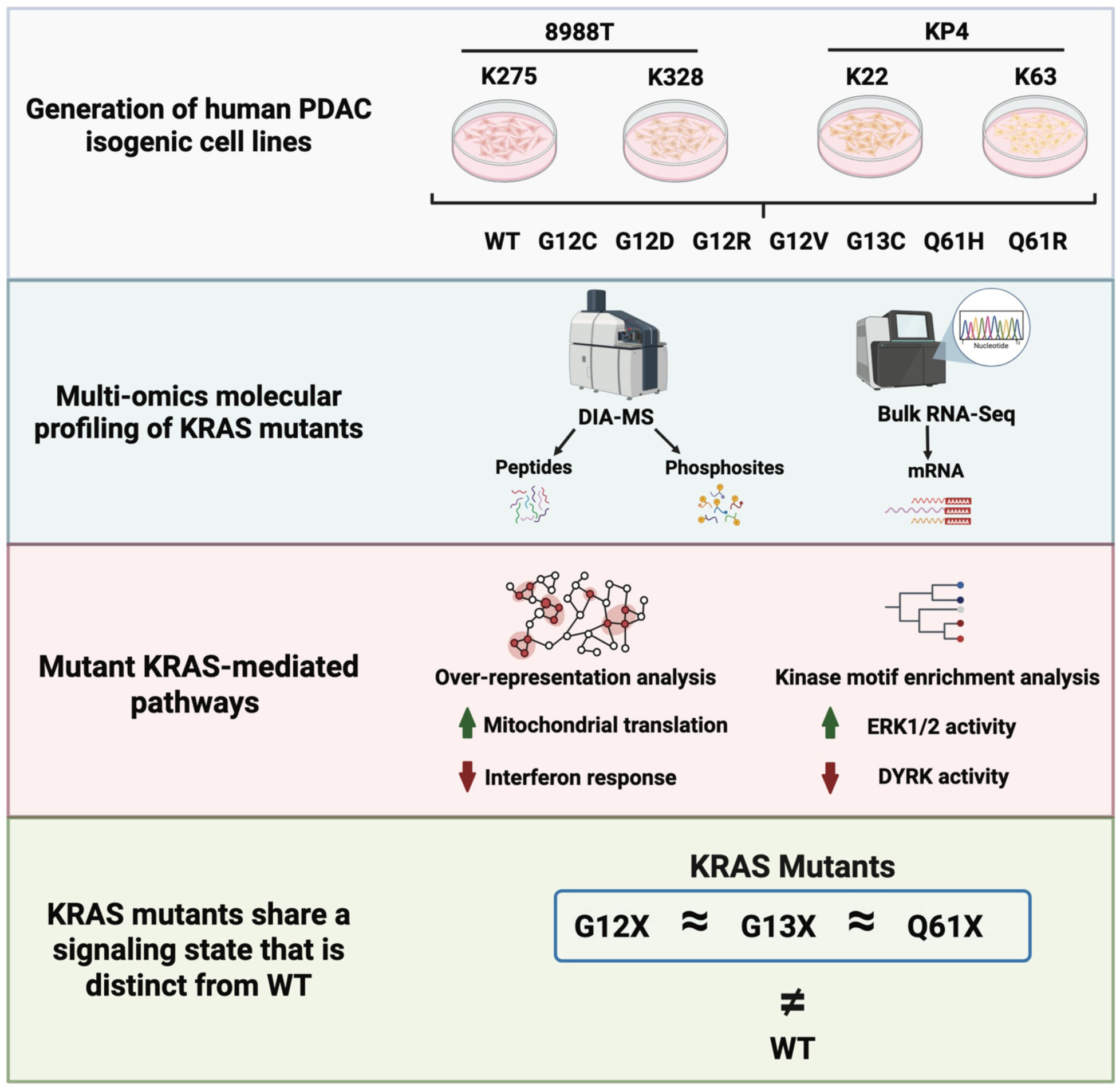
Baseline cellular state shapes KRAS mutation-dependent signaling outputs in PDAC cells. Global transcriptomic, proteomic, and phosphoproteomic (from data independent acquisition mass spectrometry (DIA-MS)) analyses of isogenic cell lines (K275, K32, K22, and K63 derived from parental 8988T and KP4 cells) reveal that *KRAS* mutant PDAC cells converge on a shared signaling state distinct from wild-type KRAS.

## SUPPLEMENTARY TABLES

**Table S1. Copy number variation analysis of *KRAS* knockout clones.** Copy number variation (CNV) analysis was performed on whole-exome sequencing data from *KRAS* knockout 8988T (K275 and K328) and KP4 (K22 and K63) clones using matched parental cell lines as controls. CNV regions are annotated with affected genes and cancer relevance based on curated databases.

**Table S2. Single nucleotide variants and indels identified in *KRAS* knockout clones.** Single nucleotide variants (SNVs) and indels were identified by whole-exome sequencing of KRAS knockout 8988T (K275 and K328) and KP4 (K22 and K63) clones using matched parental cell lines as controls. Variants shown passed filtering criteria and were functionally annotated as described in the **Materials and Methods.**

**Table S3. Protein-coding effects of somatic variants in *KRAS* knockout clones.** Protein-altering somatic variants identified in 8988T and KP4 *KRAS* knockout clones are summarized, including predicted amino acid changes and cancer relevance annotations.

**Table S4. Summary of differential expression analyses for KRAS^MUT^ versus KRAS^WT^ cell lines across transcriptomic, proteomic, and phosphoproteomic datasets.** Differential expression analyses comparing KRAS mutants (KRAS^MUT^) and matched KRAS^WT^-expressing cell lines are summarized for transcriptome, proteome, and phosphoproteome datasets across all clones. A false discovery rate (FDR) threshold of 0.05 and an absolute fold-change cutoff of 1.3 were used to determine significant differential expression per cell line.

**Table S5. Comparison of our KRAS^MUT^ transcriptomic, proteomic, and phosphoproteomic signatures with publicly available KRAS-related signatures.** Summary of overlapping features between our KRAS^MUT^ signatures and previously published KRAS-dependent signatures (Hallmark KRAS Signaling (*17*), Klomp *et al.*(*9*), Klomp *et al.*(*10*) and Kabella *et al.*(*11*)).

## REFERENCES

1. L. Rahib et al., Projecting cancer incidence and deaths to 2030: the unexpected burden of thyroid, liver, and pancreas cancers in the United States. Cancer Res 74, 2913–2921 (2014).

2. A. M. Varghese et al., Clinicogenomic landscape of pancreatic adenocarcinoma identifies KRAS mutant dosage as prognostic of overall survival. Nat Med 31, 466–477 (2025).

3. A. M. Waters, C. J. Der, KRAS: The Critical Driver and Therapeutic Target for Pancreatic Cancer. Cold Spring Harb Perspect Med 8, (2018).

4. J. C. Hunter et al., Biochemical and Structural Analysis of Common Cancer-Associated KRAS Mutations. Mol Cancer Res 13, 1325–1335 (2015).

5. D. E. Hammond et al., Differential reprogramming of isogenic colorectal cancer cells by distinct activating KRAS mutations. J Proteome Res 14, 1535–1546 (2015).

6. R. Tahir et al., Mutation-Specific and Common Phosphotyrosine Signatures of KRAS G12D and G13D Alleles. J Proteome Res 20, 670–683 (2021).

7. G. A. Hobbs et al., Atypical KRAS(G12R) Mutant Is Impaired in PI3K Signaling and Macropinocytosis in Pancreatic Cancer. Cancer Discov 10, 104–123 (2020).

8. B. Stolze, S. Reinhart, L. Bulllinger, S. Fröhling, C. Scholl, Comparative analysis of KRAS codon 12, 13, 18, 61, and 117 mutations using human MCF10A isogenic cell lines. Scientific reports 5, (2015).

9. J. A. Klomp et al., Defining the KRAS- and ERK-dependent transcriptome in KRAS-mutant cancers. Science 384, eadk0775 (2024).

10. J. E. Klomp et al., Determining the ERK-regulated phosphoproteome driving KRAS-mutant cancer. Science 384, eadk0850 (2024).

11. N. Kabella et al., Proteomic analyses identify targets, pathways, and cellular consequences of oncogenic KRAS signaling. Sci Signal 18, eadt6552 (2025).

12. M. D. Muzumdar et al., Survival of pancreatic cancer cells lacking KRAS function. Nature Communications 8, 1–19 (2017).

13. J. V. Olsen et al., Global, in vivo, and site-specific phosphorylation dynamics in signaling networks. Cell 127, 635–648 (2006).

14. Y. Liu, A. Beyer, R. Aebersold, On the Dependency of Cellular Protein Levels on mRNA Abundance. Cell 165, 535–550 (2016).

15. E. Gao et al., Data-independent acquisition-based proteome and phosphoproteome profiling across six melanoma cell lines reveals determinants of proteotypes. Mol Omics 17, 413–425 (2021).

16. J. L. Johnson et al., An atlas of substrate specificities for the human serine/threonine kinome. Nature 613, 759–766 (2023).

17. A. Liberzon et al., The Molecular Signatures Database (MSigDB) hallmark gene set collection. Cell Syst 1, 417–425 (2015).

18. C. A. Pratilas et al., ^V600E^BRAF is associated with disabled feedback inhibition of RAF&#x2013;MEK signaling and elevated transcriptional output of the pathway. Proceedings of the National Academy of Sciences 106, 4519–4524 (2009).

19. C. Vogel, E. M. Marcotte, Insights into the regulation of protein abundance from proteomic and transcriptomic analyses. Nat Rev Genet 13, 227–232 (2012).

20. M. I. Ebia, et al., Evaluating the Effect of KRAS Variants on Survival Outcomes and Therapy Response in Pancreatic Cancer. *JCO Precis Oncol* 9, e2400684 (2025).

21. C. A. McIntyre et al., Distinct clinical outcomes and biological features of specific KRAS mutants in human pancreatic cancer. Cancer Cell 42, 1614–1629 e1615 (2024).

22. S. Li, A. Balmain, C. M. Counter, A model for RAS mutation patterns in cancers: finding the sweet spot. Nat Rev Cancer 18, 767–777 (2018).

23. J. H. Cook, G. E. M. Melloni, D. C. Gulhan, P. J. Park, K. M. Haigis, The origins and genetic interactions of KRAS mutations are allele- and tissue-specific. Nature Communications 12, 1–14 (2021).

24. N. Muthalagu et al., Repression of the Type I Interferon Pathway Underlies MYC- and KRAS-Dependent Evasion of NK and B Cells in Pancreatic Ductal Adenocarcinoma. Cancer Discov 10, 872–887 (2020).

25. J. T. Park et al., Differential in vivo tumorigenicity of diverse KRAS mutations in vertebrate pancreas: A comprehensive survey. Oncogene 34, 2801–2806 (2015).

26. M. P. Zafra et al., An in vivo KRAS allelic series reveals distinct phenotypes of common oncogenic variants. Cancer Discovery 10, 1654–1671 (2020).

27. H. C. Huang et al., Kras G12C- and G12D-driven lung cancers differ in oncogenic potency, immunogenicity, and relapse after Kras inhibition in mouse models. Sci Transl Med 18, eadq6647 (2026).

28. A. M. Braxton et al., 3D genomic mapping reveals multifocality of human pancreatic precancers. Nature 629, 679–687 (2024).

29. TCGA Research Network, Integrated Genomic Characterization of Pancreatic Ductal Adenocarcinoma. Cancer Cell 32, 185–203 (2017).

30. J. Dilly et al., Mechanisms of Resistance to Oncogenic KRAS Inhibition in Pancreatic Cancer. Cancer Discov 14, 2135–2161 (2024).

31. M. M. Awad et al., Acquired Resistance to KRAS(G12C) Inhibition in Cancer. N Engl J Med 384, 2382–2393 (2021).

32. A. Sacher et al., Single-Agent Divarasib (GDC-6036) in Solid Tumors with a KRAS G12C Mutation. N Engl J Med 389, 710–721 (2023).

33. Y. Zhao et al., Diverse alterations associated with resistance to KRAS(G12C) inhibition. Nature 599, 679–683 (2021).

34. E. H. Akama-Garren et al., A Modular Assembly Platform for Rapid Generation of DNA Constructs. Sci Rep 6, 16836 (2016).

35. Y. Liu et al., Multi-omic measurements of heterogeneity in HeLa cells across laboratories. Nat Biotechnol 37, 314–322 (2019).

36. Q. Gao et al., Integrated Proteogenomic Characterization of HBV-Related Hepatocellular Carcinoma. Cell 179, 561–577.e522 (2019).

37. M. Mehnert, W. Li, C. Wu, B. Salovska, Y. Liu, Combining Rapid Data Independent Acquisition and CRISPR Gene Deletion for Studying Potential Protein Functions: A Case of HMGN1. Proteomics 19, e1800438 (2019).

38. R. Bruderer et al., Analysis of 1508 Plasma Samples by Capillary-Flow Data-Independent Acquisition Profiles Proteomics of Weight Loss and Maintenance. Mol Cell Proteomics 18, 1242–1254 (2019).

39. R. Bruderer et al., Optimization of Experimental Parameters in Data-Independent Mass Spectrometry Significantly Increases Depth and Reproducibility of Results. Mol Cell Proteomics 16, 2296–2309 (2017).

40. M. I. Love, W. Huber, S. Anders, Moderated estimation of fold change and dispersion for RNA-seq data with DESeq2. Genome Biol 15, 550 (2014).

41. D. B. Bekker-Jensen et al., Rapid and site-specific deep phosphoproteome profiling by data-independent acquisition without the need for spectral libraries. Nature communications 11, 787 (2020).

42. G. Rosenberger et al., Inference and quantification of peptidoforms in large sample cohorts by SWATH-MS. Nat Biotechnol 35, 781–788 (2017).

43. G. K. Smyth, in Bioinformatics and Computational Biology Solutions Using R and Bioconductor, R. Gentleman, V. J. Carey, W. Huber, R. A. Irizarry, S. Dudoit, Eds. (Springer New York, New York, NY, 2005), pp. 397–420.

44. H. Voss et al., HarmonizR enables data harmonization across independent proteomic datasets with appropriate handling of missing values. Nat Commun 13, 3523 (2022).

45. S. Wang et al., PTMoreR-enabled cross-species PTM mapping and comparative phosphoproteomics across mammals. Cell Reports Methods 4, (2024).

46. M. E. Ritchie et al., limma powers differential expression analyses for RNA-sequencing and microarray studies. Nucleic Acids Res 43, e47 (2015).

47. Y. Zhou et al., Metascape provides a biologist-oriented resource for the analysis of systems-level datasets. Nature communications 10, 1523 (2019).

48. Y. Perez-Riverol et al., The PRIDE database and related tools and resources in 2019: improving support for quantification data. Nucleic Acids Res 47, D442–D450 (2019).

